# Passive identification of subjective preferences towards individual items using eye-tracking in a virtual reality environment

**DOI:** 10.1101/2022.12.18.520570

**Authors:** Michal Gabay, Tom Schonberg

## Abstract

Usage of Virtual reality (VR) has been growing in many fields of research and therapy thanks to its immersive and gamified nature. Detection of the subjective experience of the users is thus essential for effective personalization of content. Eye-tracking (ET) data and specifically gaze, in two-dimensional tasks has been linked to value-based choices and emotional states. Therefore, here we aimed to develop a method for passive identification of subjective preferences based on ET data collected during a VR experience. For this purpose, we developed a naturalistic dynamic VR task where participants searched and looked at complex objects of pets and control shapes that appeared in pre-defined locations in random order. At the end of the task, participants ranked their preference, valence, and arousal of the items they saw during the task. ET data was recorded using a built-in binocular eye-tracker within the VR headset. We found that the median distance of gaze from the center of objects and the median gaze scan speed showed a significant interaction with object type (pets/shapes), as well as a significant positive relation to preference and valence rankings of pets. Our results suggest that ET could be used as a passive biomarker for detecting individual preferences and pleasantness, and in the future may enable successful personalization of VR content in real time for various applications.

## Introduction

The usage of virtual reality (VR) (Burdea and Coiffet 2003) has increased in the last decade in many research fields (Blascovich et al. 2002; Peck et al. 2013; Ossmy and Mukamel 2017; Freeman et al. 2017, 2019; Reggente et al. 2018; Hasson et al. 2019; Areces et al. 2019; Huang et al. 2021) and healthcare applications (Dahlquist et al. 2010; Cesa et al. 2013; Herrero et al. 2014; Jeffs et al. 2014; Chirico et al. 2016; Maples-Keller et al. 2017; Sacks and Axelrod 2020). Virtual reality has been shown to successfully induce affect (Herrero et al. 2014; Wu et al. 2016; Marín-Morales et al. 2018; Liao et al. 2020) more than 2D videos (Liao et al. 2020), as well as an increased sense of presence (Slater et al. 1994; Shu et al. 2018). The use of VR has been recommended for research, diagnosis and treatment of mental health disorders (Freeman et al. 2017; Rodrigues et al. 2020; Zandonai et al. 2021), for enhancing learning (Wu et al. 2020), for business and marketing (Meißner et al. 2017; Loureiro et al. 2018; Bogicevic et al. 2019; Huang et al. 2021) and even for enhancing the ecological context of MRI studies (Reggente et al. 2018; Huang et al. 2021).

In recent years personalization has become popular in various fields of research from medicine (Schleidgen et al. 2013; Schork 2015), education (Hwang et al. 2012; Xiao et al. 2017; Bernacki and Walkington 2018), through marketing (Ng and Wakenshaw 2017), to genome- and microbiome-based nutrition (Bashiardes et al. 2018). Virtual reality can also benefit from content personalization, for example to achieve a target positive emotional state in treatment of mental health disorders (Freeman et al., 2017), or an increase in learning optimization (Ravyse et al. 2017).

Thus, to allow for effective content personalization, the challenge is to be able to identify the subjective experience of users at each point in time during a VR task. To address this challenge, attempts have been made to record physiological signals. For example, to elicit and recognize affective states (valence and arousal), electroencephalography (EEG) and electrocardiography (ECG) signals were recorded during participants’ immersion in four virtual rooms, displayed via head mounted displays (HMD, Marín-Morales et al. 2018). In another study, respiration, ECG, electromyography (EMG), and electrodermal activity (EDA) signals were recorded in order to develop a biofeedback system for emotion awareness and regulation strategies in VR (Bermudez I Badia et al. 2019). However, acquisition of these physiological signals required expertise for their operation and designated equipment beyond the mere access to a VR headset, This highlights the need for subjective experience detection in VR that is within reach for common users and that does not disrupt common usage of HMDs.

The use of eye-tracking (ET) has been a promising approach in 2D studies to continuously measure subjective experiences in human-computer interfaces (Jacob and Karn 2003). It has been shown that it is possible to infer the individuals’ emotional arousal (Lanatà et al. 2013; Aracena et al. 2015) and valence (R-Tavakoli et al. 2015) by using ET while participants were observing 2D images. Gaze measures have also been linked to value-based decision-making and preferences in 2D (Shimojo et al. 2003; Krajbich et al. 2010; Krajbich and Rangel 2011; Graham and Jeffery 2012). Recent VR headsets have been pre-installed with ET devices (for example HTC-Vive https://www.vive.com/eu/product/vive-pro-eye/overview/,the Pico Neo https://www.picoxr.com/us/neo3.html, the Varjo headset https://varjo.com/products/aero/,and the Meta Quest Pro https://www.meta.com/quest/quest-pro/). Thus, ET is already accessible in advanced HMDs. One of the dominant advantages of ET in VR is experimental control on content, while mobility is preserved (Meißner et al. 2017). Examples for potential usage of ET have been demonstrated (Meißner et al. 2017) for purchase decisions in virtual supermarkets, and for user interaction with content for authentication and gaming (Khamis et al. 2018).

Most 2D ET studies focused on fixations, during which gaze is maintained on a single location; and saccades, which are ballistic eye-movements of gaze from one fixation to the other (Rayner 2009). These two ocular events characterize eye movement patterns during the majority of sedentary tasks and those that consist of viewing stationary stimuli (Lappi 2016; Kim et al. 2020). However, the eye movements that represent viewing a moving object are named smooth pursuits (Carter and Luke 2020). Considering that VR offers a naturalistic experience mimicking daily life viewing tasks, these usually include gazing at a moving object due to movement of the observer or the viewed object. Previous work with ET in VR utilized eye movements of smooth pursuits for the purpose of interaction with moving objects, where different types and features of interactions were examined (Piumsomboon et al. 2017; Khamis et al. 2018).

Here, we aimed to develop an objective and quantitative method for the identification of subjective experience measures using ET during a VR experience. Based on the above mentioned findings in 2D research, relating ET to subjective preferences and affect (measured by valence and arousal), we aimed to test whether ET measures could passively reflect individual experience in VR. We developed a novel dynamic VR task, during which participants explored a scene using head movements and gaze. The scene included pets and control shapes that moved in their location on a stationary background and participants were requested to observe the jumping stimuli. Following the viewing task, in a subsequent part of the experiment participants ranked their preference, valence, and arousal of the pets and control shapes.

We found a significant positive correlation with preference and valence rankings only for the pets category with two gaze measures: median distance of gaze from the center of the object, and the median scan speed of gaze. Both were significant while accounting for object size and percent time gazed (to account for mere exposure effects, MEE (Zajonc 1968)). These novel findings demonstrate the potential use of eye-tracking features as passive biomarkers for identifying and enhancing subjective preference and pleasantness that could be used for real-time VR content personalization. Taking into consideration the increasing availability of ET in VR HMDs, our findings have the potential to enhance optimization of different tasks in VR, such as learning, psychiatric diagnosis and treatment sessions.

## Methods

### Pre-registration

Methods for this study were pre-registered and are available in the Open Science Framework (OSF, project page: https://osf.io/2v5xn/ preregistration: https://osf.io/h9bpq).

### Participants

The participants included in this study were healthy, with intact or corrected vision between the ages of 18 and 40. The experiment exclusion criteria that were pre-defined were: participants with dizziness, pregnant women, participants with eye-glasses only for near vision correction (not contacts lenses; according to eye-tracker manufacture instructions), participants who regularly consume anti-depressants or stimulants. Participants gave their informed consent to participate in the experiment and received monetary compensation for their time of 40 NIS per our (~12$). The study was approved by the ethics committee of Tel-Aviv University.

A total of eighty-six (86) participants were recruited, out of which six participants were excluded from any analyses due to the following reasons: four participants declared eventually that they used antidepressant medication, and thus met one of the exclusion criteria. One participant did not complete the VR task due to his report of dizziness and request to discontinue the experiment. The data of one participant was not recorded due to technical issues. Additional exclusion criteria were predefined prior to data collection completion and prior to data analysis which was related to the data validity of each ranking session as detailed in the *Ranking* section below. One participant was excluded from all statistical analyses due to the ranking exclusion criteria of all ranking sessions. This resulted in a final study sample of 79 participants, (42 females) in the age of (*M* =25.28, *SD* = 4.17). Sample size rational is detailed in the *Pre-registration note* section.

### Experimental setup

The VR task was performed using a commercial HMD VR system (HTC-Vive), with screen resolution of 1080 x 1200 pixels per eye and 90 Hz refresh rate. The VR headset was adapted with a built-in binocular ET device (Tobii-installed) that provided pupil diameter data, eye gaze and head movement tracking while participants were engaged in VR. During the VR task, ET samples were recorded at a frequency of 120 Hz and a manufacturer estimated accuracy of 0.5°. The VR task was programmed and developed in C# in Unity 3D (version 2018.3) environment and ran on a dedicated computer with a Core i7-7700K CPU at 4.20 GHz, 32.0 GB RAM, an NVIDIA GeForce GTX 1080 TI GPU, and a 64-bit Windows10 operating system. The ET data acquisition was integrated in the task code and environment using Tobii’s software development kit (SDK) for Unity and edited to fit the task design and specifications.

The VR scene was based on a purchased scene from the Unity asset store, and consisted of a pastoral house (which was downloaded from: https://assetstore.unity.com/3d?category=3d&version=2018&q=ArchVizPRO%20Interior%20Vol.4&orderBy=6) and was imported to Unity and modified to fit the purpose of the task. Some of the task stimuli were also downloaded and extracted out of packages available at the Unity asset store (https://assetstore.unity.com/packages/3d/characters/animals/mammals/cute-pet-96976; https://assetstore.unity.com/packages/3d/characters/animals/fish/cute-whale-115646; https://assetstore.unity.com/packages/3d/environments/proprototype-collection-46952?category=3d&version=2018&q=PrimitivePlus&orderBy=1), and modified to fit the task purpose. Some stimuli were uniquely modeled in Blender and imported to and further designed in Unity. The participants control over the embedded task instructions, as well as the ranking selection during the ranking sessions, was performed using a trigger press on the Vive controller.

### Experimental Procedure

The general task flow comprised of two main stages beginning with a dynamic task and concluded with a ranking session (see Fig. 1).

**Fig 1.**
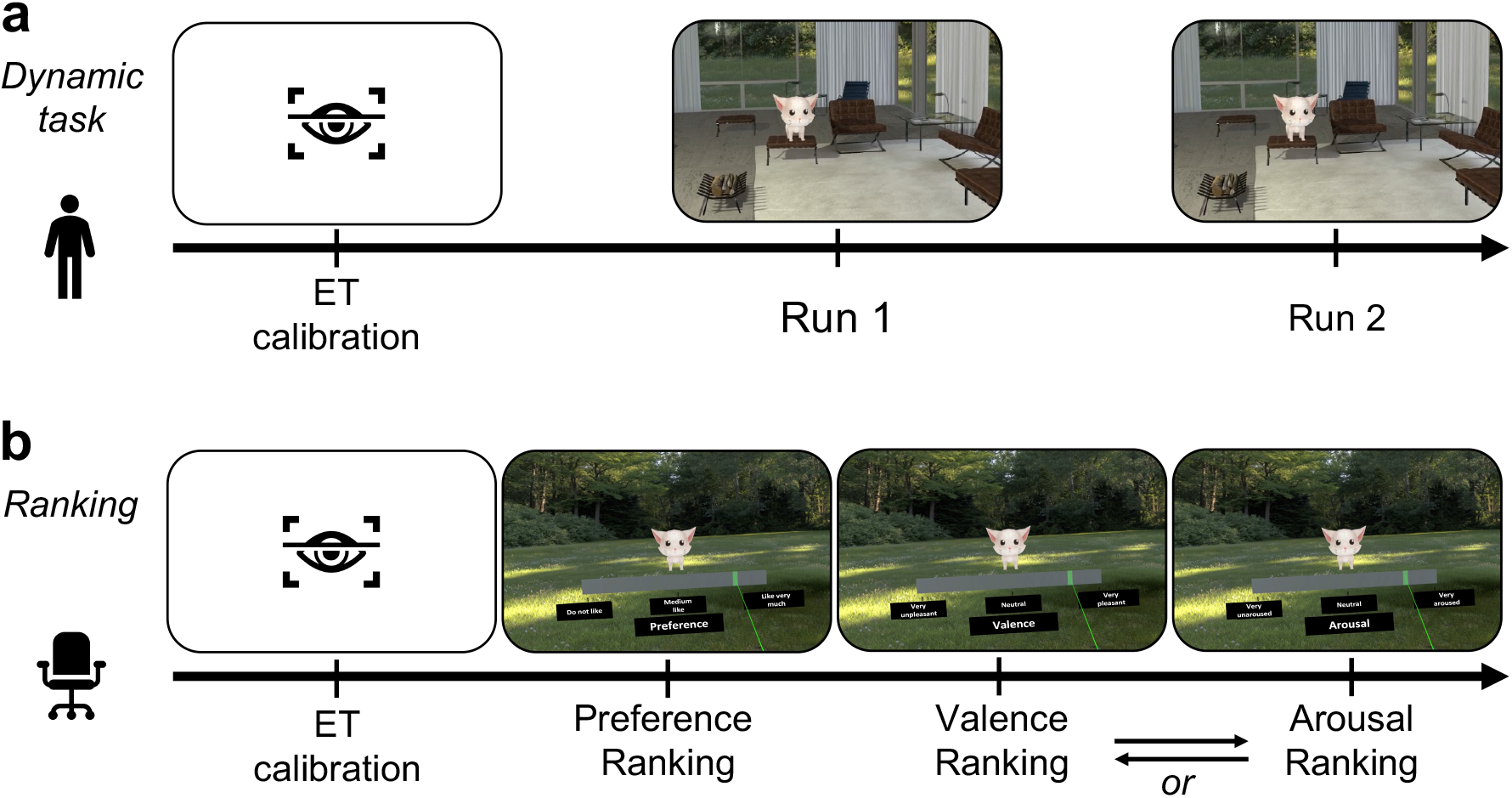
Task flow building blocks. **a:** The dynamic viewing task was conducted while standing and freely moving the head and body (no walking). The task began with an ET calibration that was followed by two runs in which the 48 stimuli appeared in random order. **b:** The ranking sessions were performed while seated and freely moving the head and hands. Following additional ET calibration all participants ranked their preference of the stimuli. Thereafter, valence and arousal ranking sessions were performed in a counter-balanced order across participants.

#### Dynamic viewing task

The placement of the HMD on participants was followed by the first ET calibration, to ensure ET signals were recorded properly while participants were engaged in the dynamic viewing task. The first part of the task was carried out while participants were standing and instructed to explore a pastoral house using body posture and head movements within the VR environment. Participants were requested to search and look at task stimuli consisting of animals (pets) and control objects (shapes) that appeared in pre-defined locations in the scene (See Fig 2.). In order to make the task closer to real life viewing conditions and to enhance participants engagement and feeling of presence, the stimuli were jumping in place up and down, and therefore were distinguishable compared to the scene furniture that were fixed. Once the objects were recognized, the participants were asked to look at them until they disappeared, and thereafter to continue and search and look at the next object, and so forth. The task included two consecutive runs, during which a total of 48 stimuli (24 pairs of animals and shapes) appeared in each run in random order (see Fig. 2). Each pet and shape pair appeared in the same location in the scene and had resembling color and size (see Fig. 2). Each object remained visible in the scene for 2 seconds from the moment it was first identified by the eye-tracker. This resulted in equal exposure times for all objects to prevent the possible confound of the mere exposure effect (Zajonc 1968, 2001; Zajonc and Markus 1982). The inter-trial interval (ITI) was at least 2 seconds, which was the time gap between the appearance of the next object after the previous object disappeared. In case the next object was not seen by the participants after 2 seconds from its appearance, a beep sound was played to inform the participants that an object is somewhere in the scene outside of their current field of view and that they needed to look for it. This was meant to enhance task engagement. The task began with 5 seconds of scene adaptation before the first stimulus was presented.

**Fig. 2:**
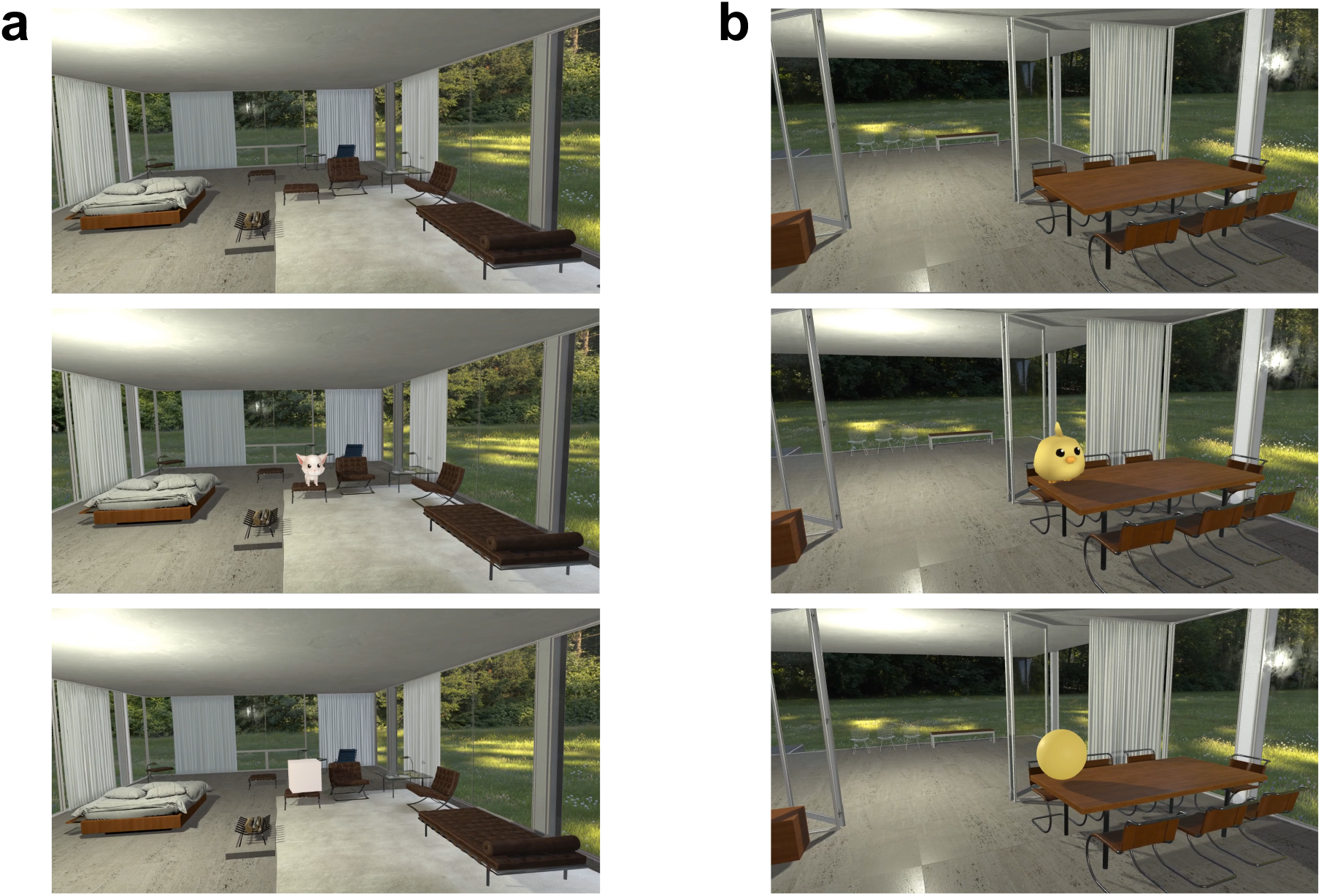
Dynamic viewing VR task setup. Pairs of pets (middle) and shapes (bottom) appeared at the same location in the scene on different trials, where the scene’s default design is shown at the top panel. **a:** a cat and a cube. **b:** a chick and a sphere. *Similar shape objects were used in the task. **Images captured from the scene are published with consent from the asset designers. (https://assetstore.unity.com/3d?category=3d&version=2018&q=ArchVizPRO%20Interior%20Vol.4&orderBy=6, https://assetstore.unity.com/packages/3d/characters/animals/mammals/cute-pet-96976

#### Ranking

Following the dynamic task, the participants ranked their preference of the stimuli they saw, and the valence and arousal levels they felt while they observed the stimuli during the task. Ranking was performed on a pseudo-continuous scale that contained unbeknownst to participants 37 bins (see Fig. 3). Therefore, the ranking score for each stimulus and for each ranking session consisted of numeric integer values in the range of 0 to 36. This part of the experiment was conducted while participants were seated and after the second ET calibration to ensure proper ET readout in the updated positioning. In the preference ranking session the participants ranked how much they liked each object on a scale ranging from “do not like” to “like very much” (see Fig. 3). In the valence ranking session the participants ranked how pleasantly they felt when they observed each object on a scale ranging from “very unpleasant” to “very pleasant”. In the arousal ranking session the participants ranked the arousal level they felt when they observed each object on a scale ranging from “very unaroused” to “very aroused”. Since preference was the main subjective measurement that we aimed to test, it was the first to be ranked for all participants. Following that, the ranking order of arousal and valence were counterbalanced across participants. During each ranking session, each stimulus was presented and ranked solely with the same neutral pastoral background, in a sequential random order with an ITI of 2 seconds. On each ranking trial, the stimulus first appeared on the screen (see Fig. 3a). Two seconds after the eye-tracker identified that participant’s gaze hit the object then a ranking scale appeared on the screen below the item (see Fig. 3b). This resulted in identical exposure time to the items to prevent the possible confound of the mere exposure effect (Zajonc 1968, 2001; Zajonc and Markus 1982) also in the ranking phase. The selection of the rank by participants was conducted in an engaging fashion, as the controller in the participants hand radiated a green “laser beam”, that colored the scale location pointed at in green (see Fig. 3b). Finally, ranking was chosen using a trigger press while the scale region of interest was pointed at and marked in green.

**Fig. 3:**
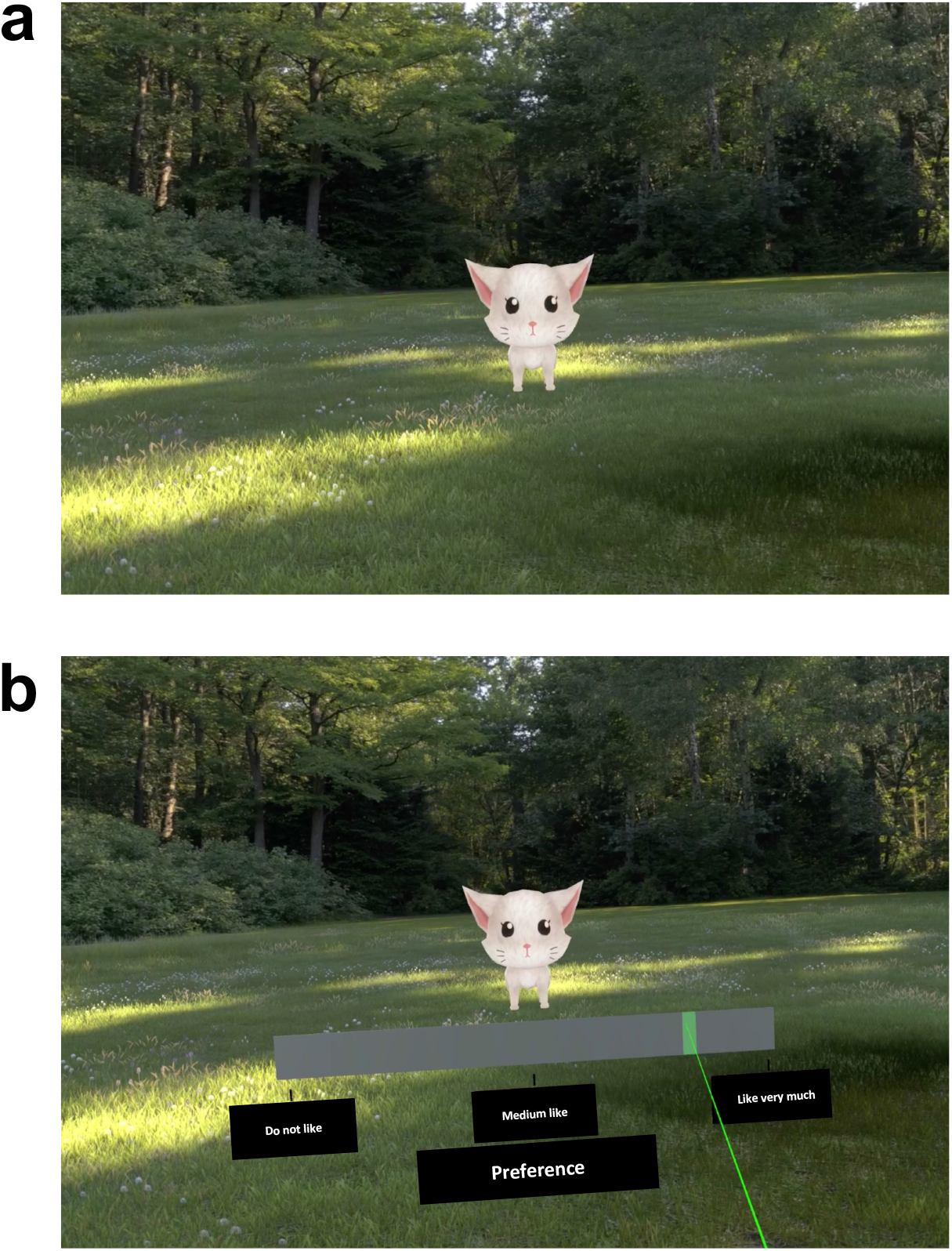
Ranking session. An example of a preference ranking trial. **a:** The ranked object, cat in this trial, first appeared by itself in the “yard” for two seconds. **b:** After two seconds since it was first viewed, the ranking scale appeared, along with its bounds and definition beneath it (the text boxes that were used in the task were in Hebrew). When the participants’ controller was pointed at a chosen region on the ranking scale it changed its color to green.

### Variables

We aimed to study eye-movements in a 3D ecological environment while viewing moving objects and thus used raw samples for the analysis of gaze behavior (Carter and Luke 2020).

Each ET data sample included measurements of: gazed ray origin (x,y,z), gazed ray direction (x,y,z), headset position origin (x,y,z), headset position direction (x,y,z), pupil diameter (mm), sample validity defined by the eye-tracker, and an arbitrary time stamp in millisecond precision. We also collected a specific time stamp readout in microseconds precision that allowed synchronization of the data with the task timeline and multiple object variables. These included: gazed object name; gazed object center position (x,y,z), for which we considered the center of the object bounding box (see Fig. 4b and c); and the gazed ray hit point (x,y,z), which was the location of collision of the gaze ray with the gazed object (see Fig. 4a). The majority of these measures were used for feature extraction as detailed in the *Feature extraction* section below. Samples validity and time stamps were used for trial validity analysis as detailed in the *Eye-tracking data pre-processing* section below.

**Fig. 4:**
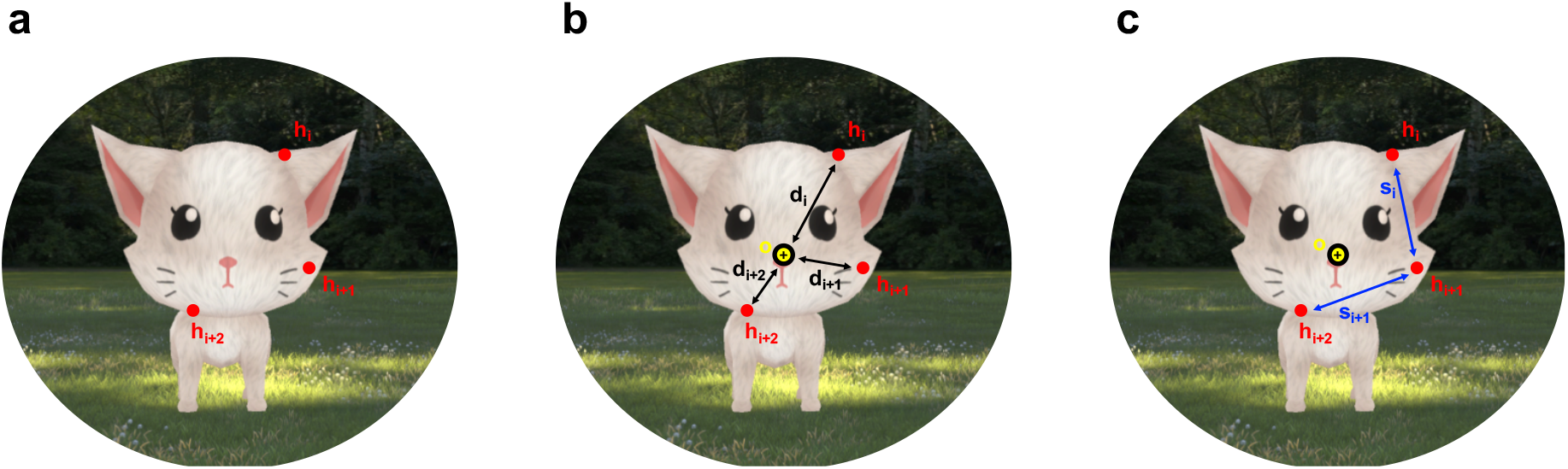
Eye-tracking features construction scheme. **a:** The location of eye gaze hit points on the cat are represented by red dots with the notation *h_i_*, *h*_*i*+1_, and *h*_*i*+2_ for the *i*th, *i*+1, and *i*+2, three time consecutive ET data samples respectively. **b:** The center of the cat’s bounding box denoted as *o* is marked by a yellow circle with a black cross in the middle, and the 3D Euclidean distances between that center and each of the gaze hit points **h_i_**, *h*_*i*+1_, and *h*_*i*+2_, are marked with black arrows and notated by *d_i_*, *d*_*i*+1_, and *d*_*i*+2_, respectively (see Equation 1 in *Feature extraction*). The Med_Dist feature is the median of these measures (see Equation 2 in *Feature extraction*). **c:** The 3D Euclidean distances between pairs of each two time consecutive gazed hit points as *h_i_* and *h*_*i*+1_ for the *i*th and the *i*+1 time points, and *h*_*i*+1_ and *h*_*i*+2_ for the *i*+1 and the *i*+2 time points, are the gaze distances per time unit, and represent the scan speed in the transition from *h*_*i*_ to *h*_*i*+1_ and from *h*_*i+1*_ to *h*_*i*+2_, are marked by blue arrows with the notation *s_i_* and *s*_*i*+1_respectively (see Equation 3 in *Feature extraction*). The Med_Scan_Speed feature is the median of these measures (see Equation 4 in *Feature extraction*). **Images captured from the task are published with consent from the assets designers. (https://assetstore.unity.com/3d?category=3d&version=2018&q=ArchVizPRO%20Interior%20Vol.4&orderBy=6, https://assetstore.unity.com/packages/3d/characters/animals/mammals/cute-pet-96976)

### Eye-tracking data pre-processing

Preprocessing of eye-tracking signals was programmed with Python 3.6 sofware (https://www.python.org/). ET samples with Nan values or non-valid samples, as defined by Tobii’s SDK, were treated as blinks. We additionally considered as blinks all samples in a 150 ms time-window before each blink onset and after each blink offset, as was done previously for 2D data in our lab (Salomon et al. 2020). All blink samples were scrubbed before the analysis. We included in the analysis only trials during which at least 50% of ET samples were valid and not Nan values, and that the stimulus gazed time was at least 50% of the 2 seconds during which it was presented from the first gaze time. Participants with less than 50% of valid trials according to these criteria were excluded from further ET analysis as in previous studies from our lab using a 2D eye-tracker (Salomon et al. 2020).

### Ranking analysis

Valid ranking data for each of the ranking sessions was defined as having a minimal standard deviation (SD) of 4 (bins) for the pets (since the power analysis registered was focused on a model for pets, and as the ranking SD of the shapes was expected to be lower than that of the pets due to their similarity to each other and relatively simple appearance). This threshold was based on two criteria that were detailed in the registration page (https://osf.io/h9bpq).

### Feature extraction

We aimed to study the link between gaze patterns on moving objects and individual experience measures. We calculated several eye-tracking features for each trial after data preprocessing. Calculated features included: percent time the object was gazed at out of the time it was presented (referred to as GazedTime), right and left pupil diameter percentiles (10th, 50th, 95th percentiles) of samples while gazing the object continuously for at least 0.5 second (referred to as Per10_PupilR/L, Per50_PupilR/L, and Per95_PupilR/L respectively); gaze shift distance percentiles (10, 50, 95) that was defined as the x,y,z Euclidian distance between the gazed hit point to the centroid of the gazed object (referred to as Min_Per_Dist, Med_Dist, and Max_Per_Dist respectively), gaze shift distance percentiles (10, 50, 95) normalized by the Euclidian distance between the gazed ray origin and the gazed object centroid (referred to as Min_Per_Norm_Obj_Dist, Med_Norm_Obj_Dist, and Max_Per_Norm_Obj_Dist respectively); blink measures included: number of blinks, total blinks duration, minimum, maximum and mean of blinks length based on raw data and according to the previous suggested constraints (Pietrock et al. 2019). We also calculated the following speed measures: the difference in time of: 1) the gaze shift distance and 2) the Euclidian distance between the participant’s head position and the gazed object position (for each the percentiles (10, 50, 95) were calculated and referred to as Min_Per_Diff_Dist, Med_diff_Dist, Min_Per_Diff_Dist, Min_Per_Diff_Obj_Dist, Med_Diff_Obj_Dist, and Max_Per_Diff_Obj_Dist respectively). For each feature the mean value for the two runs was used for further analysis. We further extracted features that represented the scan speed, as saccade velocity was found to be related to negative emotion expression (Susskind et al. 2008). Scan speed percentiles (10, 50, 95) were defined as the gaze scan Euclidian distance per time unit (referred to as Min_Per_Scan_Speed, Med_Scan_Speed, and Max_Per_Scan_Speed respectively).

An additional measure for object size was tested as a possible confound for the extracted ET features. This measure was the circumference of the bounding box that surrounded each object. This measure was referred to as DIM_SUM.

### Feature selection

The relation of all the above detailed extracted ET features to subjective experience measures was assessed using the results of a mixed linear regression model. The ranking value was the dependent variable, and the independent variables were the ET features adjusted for the object size and interacted by the viewed object type (pets vs. shapes), where participants were considered as random effects.

Since preferences were our main interest, we focused on the preference ranking regression models’ results for the purpose of feature selection. To include the possibilities of both similar and different effects according to object type, the selection criteria of features accounted for the fixed effect as well as the interaction terms in the mixed linear models. This was done by choosing the features with the smallest p value in each of these effects.

During the experiment the stimuli were presented in random order. This resulted with a diversity in participants’ body and head movements prior and during their gaze of stimuli. Consequently, the gaze ray origin and direction differed slightly from one participant to the other which affected stimuli brightness accordingly. It had been previously shown that brightness (Mathôt 2018) affects pupil diameter measures and thus we removed pupil diameter features from further analysis (nevertheless, their relation to rankings are presented in Tables 4-6 in the *Supplementary Material*).

### Selected features construction

When we ordered the extracted features in an ascending order of p-value of fixed effect term, the Med_Scan_Speed was the first from the top (i.e. lowest p value), the Med_Dist feature was the second from top (i.e. 2nd lowest p value), and Min_Per_Scan_Speed was the third from top (i.e. 3rd lowest p value). When we ordered the extracted features in an ascending order of p-value of interaction term, the Med_Scan_Speed was the first from the top, the Max_Per_Dist was the second from top, and Med_Dist was third from top. The union of the top three according to these two criteria included Med_Scan_Speed, Med_Dist, Min_Per_Scan_Speed, and Max_Per_Dist. However, since Min_Per_Scan_Speed is from the same group of features as Med_Scan_Speed, but less significant, we focused on Med_Scan_Speed out of these two. Similarly, since Max_Per_Dist is from the same group of features as Med_Dist but less significant, we focused on Med_Dist out of these two. Therefore, these two features were the focus of continued analyses according to the features selection criteria.

#### Med_Dist

The Med_Dist feature was calculated based on the gaze shift distance, noted as *d_i_* for the *i*th time point, which we defined as the 3D Euclidian distance between the gazed hit point *h_i_* for the *i*th time point and the centroid of the gazed object *o* (see Fig. 4b), where:

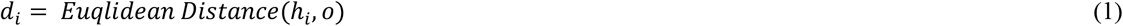

We considered the center of the objects bounding box as their 3D centroids denoted as *o*. The 50^th^ percentile of the gaze shift distances obtained in each trial was denoted as the median gaze distance, and shortly as Med_Dist:

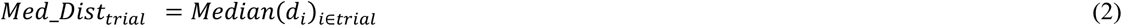

#### Med_Scan_Speed

Scan speed was defined as the gaze scan distance per time unit. Therefore, this feature was based on the distance, noted as *s_i_*, of the location of two time consequtive gazed hit points *h_i_* and *h_i+1_* for the *i*th and the *i*+1 time points (see Fig. 4c), such that:

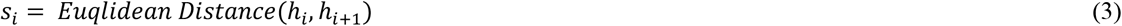

Since *s_i_* was calculated per each two time consecutive ET data samples, the corresponding time difference for all time points equaled to 8.3ms based on the 120Hz ET sampling frequency, resulting in a scaled measure for speed. The 50^th^ percentile of the scan speed per trial, is referred as Med_Scan_Speed:

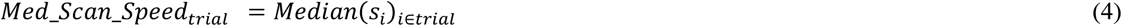

Time samples for which *d_i_* was greater than the maximal euqlidean distance obtainable for the object were considered as measurement error of the eye-tracker and excluded from further analysis.

#### Gazed Time

In addition to these gaze behavior features, we used the GazedTime feature to examine trials and participants data validity as described in the *Eye-tracking data pre-processing* section, as well as for considering the possibility of the mere exposure effect on preferences (Zajonc 1968, 2001; Zajonc and Markus 1982).

The GazedTime was defined as the percentage of time the object was gazed at out of the total time it was presented (2s).

### Statistical analyses

Statistical analysis was performed with lme4 (Bates et al. 2015) and visualizations with ggplot2 (Wickham 2016) R software (R Core Team 2022) packages. All ET features were standardized before the regression models were performed, and values greater than or smaller than 3 standard deviations from the mean were treated as outliers and were removed from the analysis. We used the standard p value of 0.05 criterion for assessing the significance of each effect. For each feature the mean value for the two runs was calculated and used for all analyses.

### Pre-registration note

This study was initiated with a pilot sample of 23 valid participants. The pilot results were used to calculate the required sample size to achieve with 80% power, a significant effect with p value < = 0.05, for the Med_Dist in the model for preference ranking. Power analysis resulted with a target sample of 53 participants with valid preference ranking, which was noted in the preregistration: https://osf.io/h9bpq. During the main study sample collection (of the n=53), three participants were excluded from the preference ranking analysis according to the ranking session exclusion criterion. Therefore, we finished the data collection with 56 participants out of which 53 with valid preference ranking.

Following completion of the main study data collection as planned and described in the pre-registration, we discovered a miscalculation that affected our eye-tracking features, including the Med_Dist feature, which we originally hypothesized upon and calculated the sample size based on.

After the calculation correction, we discovered that even though the hypothesized effect was significant in the study sample (n=53), it was not obtained in the pilot sample (n=23, see Table 1 in the *Supplementary Material*). In light of these findings, in retrospect the Med_Dist would not have been chosen as the measure to calculate the power analysis on, that determined the study sample size (details on feature selection in *Feature selection* section). However, since both samples have been collected by that point in time and we had already peeked at the data, we combined them both for reassessment and all further analysis referred in the following sections.

**Table 1:** Behavioral measures between pets and shapes. The paired difference between the pets and the shapes values of behavioral measures and their corresponding mixed linear models results: the measured difference, the t value, the degrees of freedom (df), and the p values are shown.

|  | Types difference:<br>(Pets-Shapes) | t value | df | p value |
| --- | --- | --- | --- | --- |
| Reaction Time | -0.1s | -2.8 | 23 | 0.01 |
| Gazed Time | 5.4% | 4.9 | 29.9 | 2.5e-5 |
| Preference Ranking | 24% | 6.37 | 55.8 | 1.14e-7 |
| Valence Ranking | 14% | 4 | 35.3 | 8.47e-4 |
| Arousal Ranking | 35% | 12.87 | 76.5 | 2.3e-20 |

## Results

### Valid data

For the final sample of 79 valid participants, 8.2% of the trials were excluded due to less than 50% of the objects gazed time, out of which 0.7% of the trials were excluded due to less than 50% valid or non Nan values. All participants had more than 50% valid trials, and hence were valid for ET data analysis.

### Reaction time

During the dynamic viewing task, the duration it took participants to find the animated shapes since they first appeared in the scene was (*Mean* = 1.8s, *SD* = 1.1s), which was slightly but significantly longer (in 0.1s) than the time it took to find their paired pets (*Mean* = 1.7s, *SD* = 1.2s) across all participants (*t* [23] = −2.8, *p* = 0.01, linear mixed model). The time to identify the objects was defined as the participants’ reaction time (RT), and its distribution according to object type is shown in Table 1 and Fig. 5a.

**Fig. 5:**
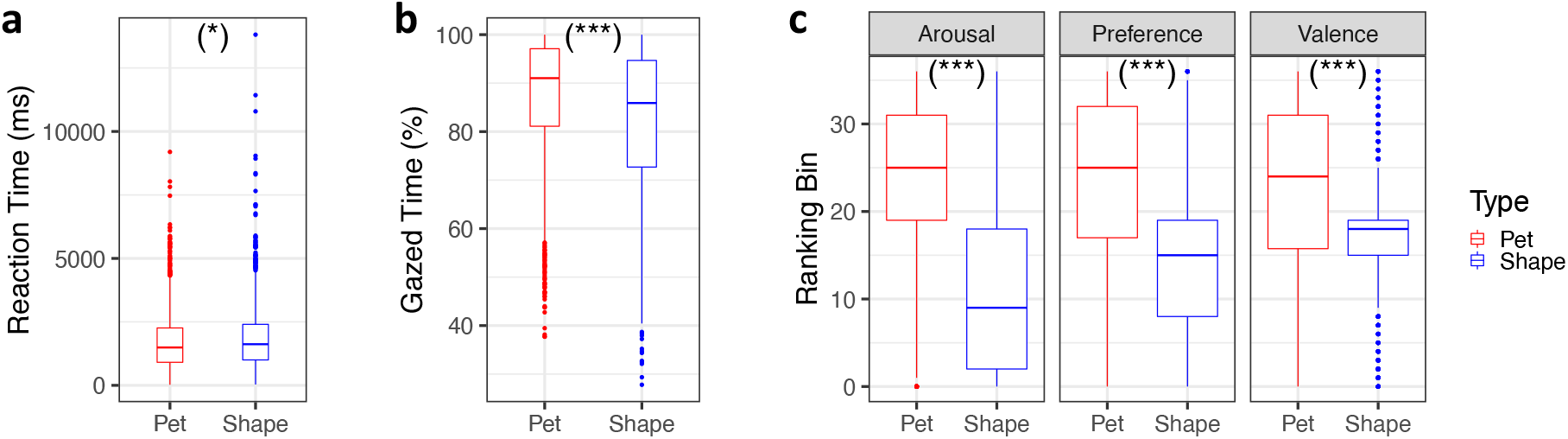
Behavioral results. Boxplots of behavioral measures according to object type (pets in red and shapes in blue). For each box, the central horizontal lines denote the median; the lower and upper borders of the box indicate the 25^th^ and the 75^th^ percentile of each distribution respectively; the whiskers denote the minimum, and maximum values that are 1.5 interquartile range from the 25^th^ and 75^th^ percentile respectively; and the dots denote values outside the whiskers range. **a:** Boxplot of the gaze reaction time (RT), that was defined as the time of the first gaze point on the object since the time it appeared in the scene. **b:** Boxplot of the percent time gaze out of 2 seconds each object was presented since it was first observed. **c:** Boxplots of the objects’ rankings of arousal, preference, and valence. (Significance is noted for the test for the difference in each measure for pairs of pets and shapes using a linear mixed model, Bonferroni corrected for 3 ranking tests, where p < 0.001 ‘***’, p < 0.01 ‘**’, p < 0.05’*’, p < 0.1 ‘.’).

### Gazed time

Participants were instructed to look at the appearing stimuli from the moment they were first observed until they disappeared. Nevertheless, we found that the percent gazed time was significantly longer by 5.4% for pets (*M* = 87.25%, *SD* = 12.38%) than for their paired control shapes (*M* = 82.37%, *SD* = 15.19%) across all participants (*t*[29.9] = 4.9, *p* = 2.5e-5, linear mixed model, Table 1 and Fig. 5b). Therefore, to account for possible mere exposure effects (MEE, (Zajonc 1968, 2001; Zajonc and Markus 1982)) we added the percent of gazed time as an additional independent variable to all subsequent ranking models.

### Ranking

The ranking sessions validity criterion yielded 75, 77, and 66 participants with valid ranking data for the preference, valence, and arousal rankings respectively.

As expected, the preference, valence, and arousal rankings consisted of significantly higher values for pets than for their paired control shapes (Table 1 and Fig. 5c). After Bonferroni correction for 3 tests in 3 mixed linear regressions we found a significant difference of 24% of the scale range in the pets preference ranking compared to shapes (*t*[55.8] = 6.37, *p* = 1.14e-07); a significant difference of 14% of the scale range in the pets valence ranking compared to shapes (*t*[35.3] = 4, *p* = 8.47e-4); and a significant difference of 35% of the scale range in the pets arousal ranking compared to shapes (*t*[76.5] = 12.87, *p* = 2.3e-20). The gazed time we added to the model did not have a significant effect on any of the rankings (*p* >0.4 in all models for the effect of gazed time).

### Feature selection

As explained above in the *feature selection* section the features were sorted in ascending order according to the fixed effect p value and interaction p value. Table 2, is sorted according to the fixed effect parameter and shows the Med_Scan_Speed was the first, and the Med_Dist was the second from the top.

**Table 2:** Ranking of eye tracking features in relation to preference ranking. For each features’ mixed linear model: the fixed effect estimate, the fixed effect p value, the interaction estimate, and the interaction p value are shown. Features are sorted in ascending order according to the fixed effect p value, that was one of the selection criteria for our main features * Features relation to valence and arousal rankings are shown in Tables 2 and 3 respectively in the *Supplementary Material*).

|  | Feature | Fixed effect estimate | Fixed effect P value | Interaction estimate | Interaction P value |
| --- | --- | --- | --- | --- | --- |
| 1 | **Med_Scan_Speed** | 1.80E-01 | 3.11E-13 | -1.88E-01 | 4.71E-11 |
| 2 | **Med_Dist** | 1.83E-01 | 2.16E-12 | -1.87E-01 | 3.65E-09 |
| 3 | Min Per Scan Speed | 1.42E-01 | 7.50E-11 | -1.58E-01 | 4.07E-08 |
| 4 | Max Per Dist | 1.61E-01 | 1.12E-10 | -1.96E-01 | 3.88E-10 |
| 5 | Med Diff Obj Dist | 1.26E-01 | 5.85E-10 | -1.12E-01 | 8.98E-05 |
| 6 | Med_diff Dist | 1.22E-01 | 1.10E-07 | -1.60E-01 | 8.54E-09 |
| 7 | Max Per Diff Dist | 1.04E-01 | 7.97E-07 | -1.52E-01 | 4.04E-08 |
| 8 | Max Per Scan Speed | 9.14E-02 | 2.63E-06 | -1.10E-01 | 5.55E-05 |
| 9 | Max Per Diff Obj Dist | 7.93E-02 | 1.21E-05 | -9.22E-02 | 6.34E-04 |
| 10 | Min Per Diff Dist | 9.40E-02 | 1.49E-04 | -1.27E-01 | 1.76E-05 |
| 11 | Max Per Norm Obj Dist | -7.09E-02 | 3.19E-04 | 7.40E-02 | 6.94E-03 |
| 12 | Min Per Dist | 8.32E-02 | 1.00E-03 | -7.78E-02 | 1.12E-02 |
| 13 | Blinks Sum Constrained | 6.42E-02 | 3.02E-03 | -9.86E-02 | 4.18E-04 |
| 14 | Blinks_Sum | 6.02E-02 | 4.70E-03 | -9.08E-02 | 1.29E-03 |
| 15 | Blinks_Mean | 5.70E-02 | 8.98E-03 | -8.48E-02 | 2.15E-03 |
| 16 | Blinks_Mean Constrained | 5.36E-02 | 1.62E-02 | -8.48E-02 | 1.97E-03 |
| 17 | Min Per Norm Obj Dist | -4.70E-02 | 1.75E-02 | 6.72E-02 | 1.82E-02 |
| 18 | GazedTime | -5.67E-02 | 1.83E-02 | 5.90E-02 | 4.70E-02 |
| 19 | Blinks Num Constrained | 4.54E-02 | 1.93E-02 | -6.84E-02 | 1.57E-02 |
| 20 | Blinks Max Constrained | 4.96E-02 | 2.41E-02 | -8.48E-02 | 2.13E-03 |
| 21 | Blinks_Num | 4.26E-02 | 3.01E-02 | -5.72E-02 | 4.18E-02 |
| 22 | Blinks_Min Constrained | 4.41E-02 | 3.61E-02 | -6.70E-02 | 1.22E-02 |
| 23 | Blinks_Max | 4.58E-02 | 3.91E-02 | -7.31E-02 | 8.85E-03 |
| 24 | Med Norm Obj Dist | -3.26E-02 | 8.30E-02 | 4.97E-02 | 7.61E-02 |
| 25 | Blinks_Min | 3.62E-02 | 8.33E-02 | -4.92E-02 | 6.70E-02 |
| 26 | Min Per Diff Obj Dist | 2.42E-02 | 2.21E-01 | -1.57E-03 | 9.56E-01 |

### Selected features

The median distance of gaze, referred to as Med_Dist, was significantly higher for pets relative to their paired control shapes across all participants (*t*[22] = 2.67, *p* = 0.028, mixed linear regression, adjusted for gazed time and object size, Fig. 6). This feature represents a measure for how far the participants scan the stimuli away from their center. Therefore, this difference between the pets and shapes could reflect or be influenced by the greater complexity and points of interest the pets have in regions proximal to bounds relative to the shapes. The gazed time in this model had a trending effect on the Med_Dist values (*t*[1774] = −2, *p* =0.08, mixed linear regression), but still Med_Dist is significant above and beyond gazed time.

**Fig. 6:**
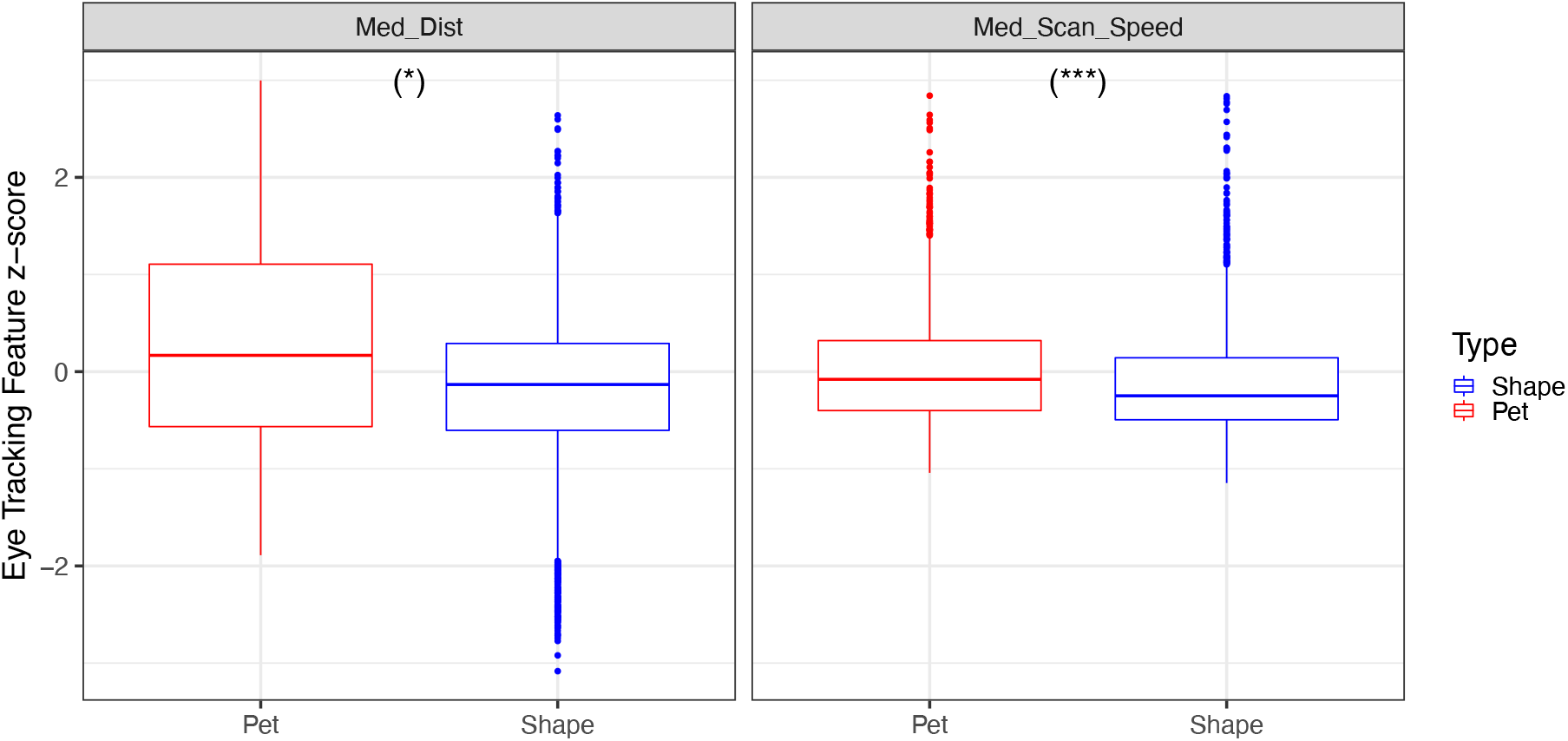
Selected features distribution according to object type. Boxplots of eye-tracking features according to object type (pets in red and shapes in blue). For each box, the central horizontal lines denote the median; the lower and upper borders of the box indicate the 25^th^ and the 75^th^ percentile of each distribution respectively; the whiskers denote the minimum, and maximum values that are 1.5 interquartile range from the 25^th^ and 75^th^ percentile respectively; and the dots denote values outside the whiskers range. Significance is noted for the tests for the difference in each measure for pairs of pets and shapes using a linear mixed model adjusted for the gazed time and objects size. Bonferroni correction for the 2 features found a significant paired difference for both Med_Dist and Med_Scan_Speed features (where p < 0.001 ‘***’, p < 0.01 ‘**’, p < 0.05‘*’, p < 0.1 ‘.’).

A similar relation was found for the median scan speed feature, referred to as Med_Scan_Speed, where significantly higher values for pets were obtained relative to their paired control shapes across all participants (*t*[54.2] = 5.47, *p* = 2.3e-06, mixed linear regression, adjusted for gazed time and objects size, Fig. 6). Unlike for Med_Dist, the paired difference in Med_Scan_Speed was significantly negatively influenced by the gazed time (*t*[21.4] = −4.1, *p* = 8.6e-4, mixed linear regression). This could result from an attempt to compensate higher scan speed for less time of viewing an object. These ET results were Bonferroni corrected for the 2 features.

### Correlation of eye tracking and subjective experience measures

#### Med_Dist relation to subjective rankings

We tested the relation of Med_Dist to ranking using a mixed linear model adjusted for objects size and gazed time to control for the possible mere exposure effect (Zajonc 1968, 2001; Zajonc and Markus 1982). The model included object type interaction, and the participants were modeled as random effects (Bonferroni corrected for 3*32 tests, i.e., 96). A significant interaction of Med_Dist and type was found for preference (*t*[106.4] = −4.9, *p* = 2.6e-4, Fig. 1a in *Supplementary Material*) and valence and (*t*[222.2] = −6.2, *p* = 2e-7, Fig. 1b in *Supplementary Material*). In addition, we found a significant positive relation between Med_Dist and preference of pets (*beta* = 0.18, *t*[70.4] = 4.6, *p* = 0.0016) and valence rankings of pets (*beta* = 0.25, *t*[105] = 7.4, *p* = 3.3e-9). This implies that the furthest away from the center of a pet the participants fixated upon, the more they liked it, and the more pleasant they felt while looking at it. This effect was not obtained for the shapes (p > 0.87 for both preference and valence).

We additionally tested the Med_Dist relation to arousal rankings. We did not find any significant interactions (*t*[606] = 1.6, *p* = 1). However, a significant main effect of Med_Dist was found in relation to arousal rankings (*beta* = −0.13, *t*[581] = −5.5, *p* = 5.3e-6, Fig. 1c in *Supplementary Material*) suggesting that across all items the furthest away participants looked from the center the less they were aroused.

The gazed time had no effect on the rankings in these models (*p* = 1, for both main effect and interaction with type in all corresponding models).

#### Med_Scan_Speed relation to subjective rankings

The relation between Med_Scan_Speed and subjective rankings was tested using a linear mixed models adjusted for objects size and gazed time, including object type interactions, where participants were considered as random effects (Bonferroni corrected for 3*32 tests).

A significant interaction of Med_Scan_Speed and type was found for preference (*t*[83] = −5.6, *p* = 2.88e-5, Fig. 7a) and valence (*t*[75.5] = −6.6, *p* = 5.3e-7, Fig. 7b). In addition, we found a significant positive relation between Med_Scan_Speed and preference of pets (*beta* = 0.19, *t*[68.3] = 6.2, *p* = 4.1e-6) and valence of pets (*beta* = 0.28, *t*[66.7] = 6.9, *p* = 2.1e-7), while no relation was obtained for the shapes type (*p* = 1 for both). These results suggest that the faster participants scanned the pets with their gaze, the more they liked them, and the more pleasant they felt while looking at them.

**Fig. 7:**
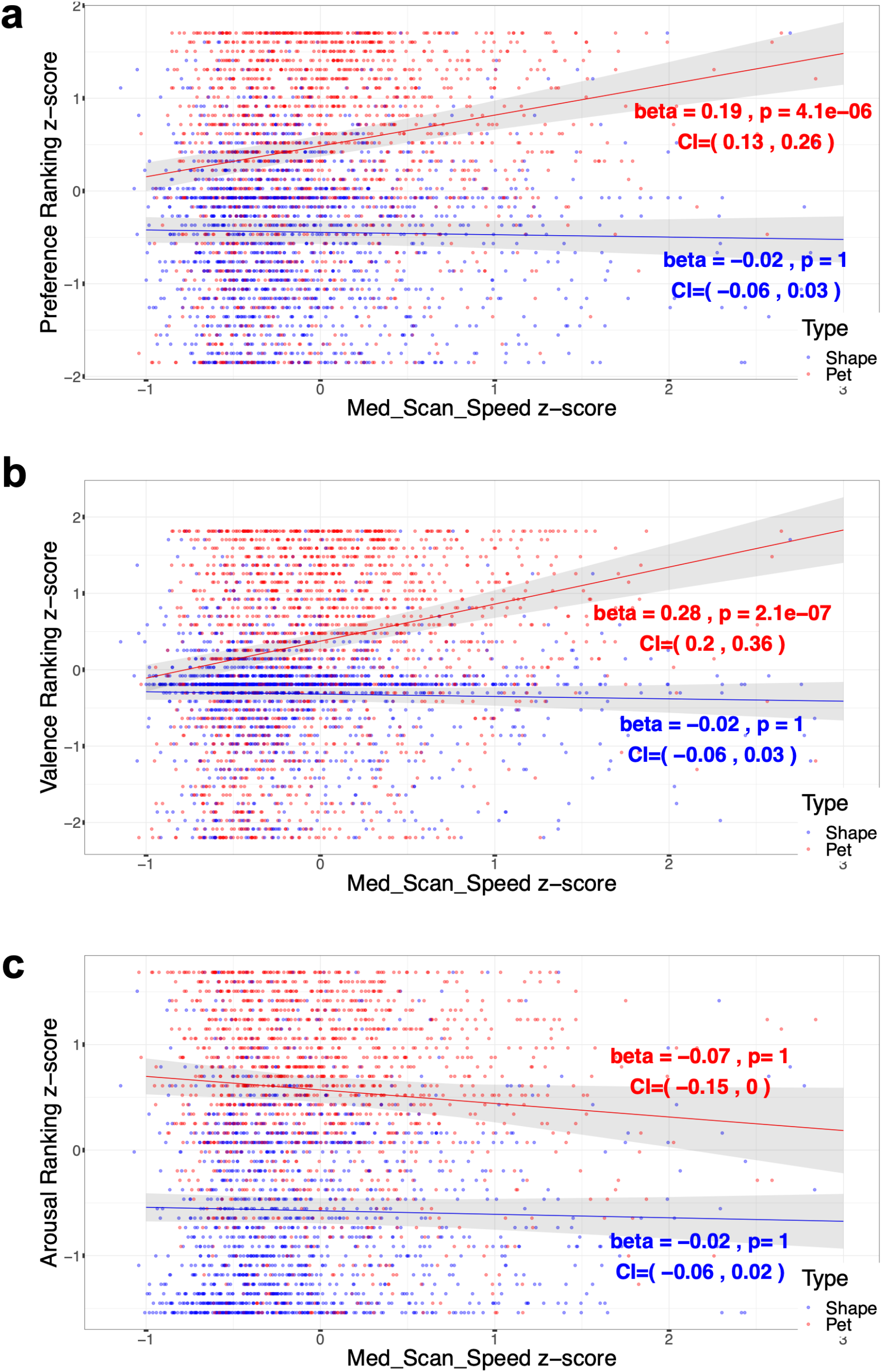
Med_Scan_Speed relation to rankings. Scatter plots of the subjective rankings as a function of Med_Scan_Speed per participant per object, and the linear models predicted values of rankings as a function of Med_Scan_Speed for each object type (pets in red and shapes in blue; fitted lines are surrounded by gray polygons of the 95% confidence interval). **a:** Preference rankings as a function of Med_Scan_Speed. A significant interaction of Type was obtained (beta = −0.2, t[83] = −5.6, p = 2.88e-5), CI = (−0.28,−0.14)) as described by a significant positive slope only for the pets in the plot. **b:** Valence ranking as a function of Med_Scan_Speed. A significant interaction of Type was obtained (beta = −0.3, t[75.5] = −6.6, p = 5.3e-7, CI = (−0.39,−0.21)) as described by a significant positive slope only for the pets in the plot. **c:** Arousal ranking as a function of Med_Scan_Speed. Type interaction was not found for arousal ranking (beta = 0.05, t[53.3] = 1.2, p = 1, CI = (−0.03,0.14)) as described with two non-significant slopes for both types in the plot. (All variables are z-scored, and significance is noted for the Med_Scan_Speed estimate in a mixed multilinear model adjusted for objects dimension and percent time gaze for each type, where p < 0.001 ‘***’, p < 0.01 ‘**’, p < 0.05‘*’, p < 0.1 ‘.’; all tests are Bonferroni corrected for 3*32, i.e., ranking times the number of features = 96 tests).

As for the arousal rankings (Fig. 7c), the Med_Scan_Speed feature did not have a significant interaction with type (*t*[53.3] = 1.2, *p* = 1) nor a significant main effect (*t*[53] = −1.9, *p* = 1).

The gazed time had no effect on the rankings in these models (*p* = 1 for both main effect and interaction with type in all these models).

#### The relation of ET to the paired ranking difference

Our assumption in pairing a control shape to each pet was that they will be ranked differently, despite their similar color, size, and position in the scene. As detailed above, we found a significant positive relation between Med_Dist and Med_Scan_Speed to pets’ preference and valence rankings accompanied by a significant interaction with shapes, and a significant paired difference of these rankings and ET measures. Therefore, even though these ET features had no effect on the shapes valence and preference rankings, we further tested whether the paired difference in the ET measures is related to the paired ranking difference for each pair of pet and its control shape. We tested this with a mixed linear regression fitted to the paired ranking difference, this time in a combined model with both the paired difference in Med_Dist and Med_Scan_Speed as independent variables adjusted for the paired difference in size and gazed time. After Bonferroni correction for the two rankings of preference and valence, we found that the paired difference in the Med_Scan_Speed was positively related to the paired difference in preference ranking (*beta* = 0.12, *t*[15.3] = 3.6, *p* = 0.005) and valence rankings (*beta* = 0.14, *t*[15.4] = 3.8,*p* = 0.003). Nevertheless, the paired difference in Med_Dist was not correlated with the paired difference in ranking (*t*[16.7] = −1.2, *p* = 0.5) and (*t*[17.5] = −1.3, *p* = 0.4) for preference and valence ranking respectively. No relation was found for both paired differences in gazed time and objects size to the paired ranking difference (*p*>0.27 for all). These results suggest that the higher the difference in the characteristic Med_Scan_Speed of pets relative to their control shapes, the greater the subjective difference in their preference and in the valence that participants felt while looking at them.

## Discussion

Virtual reality has been gaining popularity in research (Blascovich et al. 2002; Peck et al. 2013; Ossmy and Mukamel 2017; Freeman et al. 2017, 2019; Reggente et al. 2018; Hasson et al. 2019; Areces et al. 2019; Huang et al. 2021) and therapeutics (Dahlquist et al. 2010; Cesa et al. 2013; Herrero et al. 2014; Jeffs et al. 2014; Chirico et al. 2016; Maples-Keller et al. 2017; Sacks and Axelrod 2020) due to its immersive gamified nature. Simple personalization of presented content is highly needed to increase its effectiveness. In this study we aimed to address this personalization challenge by using eye tracking of moving individual objects. We devised a novel task whereby we presented participants with individual moving objects and analyzed eye gaze data. We found that the larger the scanned distance from the center of an object (named Med_Dist) and the scan speed (Med_Scan_Speed) of that object, the greater the participants’ preference of this object. The gazing on individual items and ranking of items were performed in separate phases of the experiment. This suggests that gaze tracking while passively viewing items can serve as a marker for preference without the need to probe it directly. We compared two stimuli categories of pets and of shapes, controlling for color, size, and scene position of the pets compared to shapes. We tested multiple gaze features, and following a feature selection process, we focused on two features, median distance and median scan speed. We found a positive correlation between these. ET features and the subjective rankings of preferences as well as the valence of viewed pets. These results suggest that ET measures extracted during passive viewing of individual moving objects during a VR experience could be used as biomarkers of subjective preference and pleasantness.

Findings relating gaze and preferences have been studied in tasks with 2D images of faces and shapes (Shimojo et al. 2003), snacks (Krajbich et al. 2010; Krajbich and Rangel 2011) or food items (Graham and Jeffery 2012). During these tasks participants were requested to choose the one they preferred out of two (Shimojo et al. 2003; Krajbich et al. 2010) or three (Krajbich and Rangel 2011) presented items. Alternatively they were requested to indicate it they would buy an item in a simulated food purchase (Graham and Jeffery 2012). The common result in these studies was that the accumulated amount of time gazed, also referred to as dwell time, was positively related to preference or choice (Shimojo et al. 2003; Krajbich et al. 2010; Krajbich and Rangel 2011; Graham and Jeffery 2012). In addition, with the development of an extended version of the drift diffusion model (DDM), a more complex relationship was found between fixation patterns and choices, as well as several fixation-driven decision biases in binary (Krajbich et al. 2010) and trinary (Krajbich and Rangel 2011) value based choice tasks. Additional ET features in these studies were the item’s identity and the dwell time at the first and last time points (Krajbich et al. 2010; Krajbich and Rangel 2011). Thus, while prior work examined gaze and preferences during active choice, in the current study we set to investigate passive gaze time series patterns of individual items with the purpose of detecting subjective experience without choice or valuation elicitation. Therefore, our task was designed to study passive viewing we often encounter in daily life, which includes data gathering on objects presented one at a time, without necessarily applying a subsequent choice or action.

Previous studies employing 2D stimuli found that the amount of time an object was gazed at influenced preference (the mere exposure effect, MEE, (Zajonc 1968, 2001; Zajonc and Markus 1982; Shimojo et al. 2003). With the attempt to control for this potential confound, all participants were instructed to look at each stimulus from the moment it was first noticed and until it disappeared. Still, a small but significant difference was obtained between the percent time the pets were gazed at, relative to their control shapes. To account for MEE, we added this variable to our models and found it had no relation to preference or any of the other ranking measures.

The positive correlation between both Med_Dist and Med_Scan_Speed features to subjective rankings was demonstrated from multiple perspectives. These two ET features obtained higher values for the pets relative to their control shapes. Similarly, the pets were also ranked higher than shapes for the measures of preference, valence, and arousal. A possible alternative explanation for this general difference in ET features between the two object types could have been influenced by the greater complexity and salient details of the pets such as eyes, mouth, ears, and tail, whereas the shapes lacked these properties. Participants could have been attracted to observe these features that are close to the pets’ perimeter, and thus derive their greater values of Med_Dist. Furthermore, a possible attempt to capture the additional details in the pets could have led to their obtained higher Med_Scan_Speed. This idea is in line with the negative relation found between Med_Scan_Speed and percent gazed time, that could point at an attempt to compensate for the shorter time gazed while collecting the same amount of data on the viewed object. We additionally found a positive relation between the difference in Med_Scan_Speed of pairs of pets and their control shapes and the corresponding difference in their preference and valence rankings. The percent gazed time had no effect on this paired difference in valence and preference rankings. Taken together, these results suggest that the gazed time relation to Med_Scan_Speed describes a difference between the categories that is unrelated to the difference in their liking and pleasantness.

A positive relation of these ET features to preference and valence rankings was found within the pets category alone, suggesting a characteristic difference in gaze patterns when looking at shapes vs. pets. A possible explanation for this interaction with object category is the greater number of salient details in pets, that could have contributed to a greater diversity in gaze behavior in this category compared to their control simple shapes. In a study that classified images by modeling scan paths with variational hidden Markov models (HMMs), a positive correlation was obtained between the classification accuracy and the amount of salient objects in the image (Coutrot et al. 2018). This suggests that objects complexity allows for better inference of their content based on gaze behavior. Taken together with the findings in our study, it seems that the greater the complexity or salient items 2D or 3D objects have, the greater is the ability to deduce their content, preference, and possibly other traits, based on ET data.

Object size was previously tested but not found to be related to affective preferences when it was examined in a controlled fashion (Seamon et al. 1997). Other studies found that preference of different stimuli categories was affected by different visual attributes, although size was not tested (Oren et al. 2020). Therefore, since both the gaze distance (Med_Dist) from the center of the object, and the Med_Scan_Speed of the object could have been influenced by the object size, we included the objects’ circumference as a measure for its size to all our mixed linear models for rankings prediction. We found that both ET features of Med_Dist and Med_Scan_Speed were positively related to pets’ preference and valence ranking adjusted for stimuli size. This result suggests that extended gaze at peripheral points and higher scan speed characterize the exploration pattern gaze at pets, regardless of their size. Furthermore, this gaze pattern indicates higher subjective preference or valence. The pets in this study may be considered as an example for complex objects, when compared to their control shapes.

In contrary to the positive relation found for the Med_Dist feature to pets preference and valence rankings, a negative relation was obtained for both the pets and shapes arousal rankings. This result was initially surprising. Therefore, we examined in depth the identity of the objects that were mostly ranked to induce high arousal levels. We found that they were all also ranked with low valence (and low preference for the pets in this group). Hence high arousal stimuli generated the opposite trend for valence, and preference in the pets group (for details see section *S5* in *Supplementary Material*). Related work, aimed to investigate the relation between ET features and emotional states, was done on 2D images using the International Affective Picture System (IAPS) (Lanatà et al. 2013; R-Tavakoli et al. 2015; Aracena et al. 2015), which are standardized emotion inducing images (Lang, P. J., Bradley, M. M., & Cuthbert 1997). Those studies aimed to classify one dimension of affect as arousal levels (Lanatà et al. 2013), valence (R-Tavakoli et al. 2015), or positivity level (Aracena et al. 2015) and used a subsample of the IAPS images accordingly. In contrast, we studied the relation of ET features to two affect dimensions on 3D objects. The affect rankings in studies that used the IAPS over the years are robust compared to introducing and ranking stimuli for the first time as done in our study, which is one of its drawbacks. It seems that standardizing 3D emotional objects to be used in VR studies of affect could significantly contribute to the field as the IAPS has been for 2D images. Our study included only two object types of pets and shapes as our design aimed to keep the task at a reasonable time frame for wearing the VR headset. Thus, future work will be required to generalize our findings to other types of objects, environments, and scenarios.

Most ET studies thus far were done on sedentary tasks and in front of a 2D screen (Lappi 2016) sometimes with the addition of a chinrest, to ensure all participants look at the screen from the same point of view. However, in many natural gazing tasks we encounter in our daily lives, we look at objects from different points of view and even during movement. In our study, even though all stimuli appeared at the same locations for all participants, since they appeared in a random and therefore different order they were observed from possibly slightly different angles, points of views and perspectives due to participants’ individual body, head, and eye movements prior to the stimuli being observed. As a result, participants might have not all looked at the exact same coordinates on all stimuli. Thus, extracting global features for spatial dynamics of the eye scan of objects as we performed, is more compatible than a coordinate-wise fixation location analysis usually done in 2D studies. Notwithstanding that this adds variance to the data, it also allows experimenting in a naturalistic fashion. Therefore, this current study provides novel analysis and insights of gaze behavior in a naturalistic engaging environment. Hence, it could also be applicable to studies with eye-tracker built-in glasses as it allows free head and body movements as in VR systems.

Passively identifying the individual preference or pleasantness of various objects or figures in a computerized scene, simulation, or training session, will allow real time updating of the content presented in the direction of the preferred content and will increase the participant enjoyment and engagement. These were previously shown to enhance learning in serious games (Ravyse et al. 2017). Therefore, using the methodology presented in the current work may contribute to optimization of a computerized learning or training tasks to enhance learning. In an unpublished study, we established a closed loop system for this task in our lab that identifies the individual preference of stimuli based on the selected ET features and updated the task in real time accordingly. In one application, the size of the following presented stimulus was changed according to the preference of its previous one. In another application, a group of stimuli was presented according to the preference inference of an initial group. When the entire experience was ranked, greater enjoyment was found for participants who were presented with stimuli with higher preference, than those with lower (data not shown). This may apply to different regimes as: elementary or high school and other education systems; skills training tasks as for surgeons virtual training; factory employee machinery training; different private training software for language, programing, talking in front of an audience; or any knowledge transferred via software in a 2D screen or VR headset format. These principles may be also adopted for gaming, where enhancing the individual enjoyment and engagement, may virtually contribute to improving the game. In addition, preference identification may be used for personalization of in-game purchases.

For therapeutic purposes, an example could be with the attempt to achieve a target positive emotional state in treatment of depression, anxiety, or for example as a distraction for when removing bandages after burn injuries (Jeffs et al. 2014). Personalization of the presented virtual content could be done according to the identification of the individual preference and feeling of pleasantness of the presented objects, surrounding people, scenery design, trees, etc. and increase the therapy’s impact. In addition, user preference calculation in the framework of psychotherapy software may be used to evaluate the progress and effect of psychotherapy during the execution of the software, or over multiple different executions/sessions, by measuring the difference in user preference to the presented objects throughout the session(s). These are just a few examples for utilizing our findings in future real-time updated applications, that are likely to grow with the technological and scientific developments in the coming years.

To conclude, the concept of VR content updating based on subjective internal states has been studied using physiological signals (EEG and ECG (Marín-Morales et al. 2018), respiration, ECG, EMG, and EDA (Bermudez I Badia et al. 2019)). However, these necessitated additional equipment and expertise. Here, we show that using a built-in eye-tracker and unique analyses we could identify preferences and valence. Finally, the use of ET in human-computer interfaces has been noted as a promising approach either alone or combined with inputs from other sensors or devices (Jacob and Karn 2003). The utilization of the immersive naturalistic nature of VR is increasing in various research fields and applications. Therefore, our findings suggest that ET measures could passively reflect an individual internal preference and state. In turn, these could be used for content personalization, either with or without additional physiological signals, via real time updating of VR tasks for research, learning, or therapeutic purposes.

## Funding

Funding for this research was provided by the Minducate Science of Learning Research and Innovation Center of the Sagol School of Neuroscience, Tel Aviv University and The Zimin Institute for Engineering Solutions Advancing Better Lives.

We thank Dr. Jeanette Mumford for statistical consulting, Tom Salomon for helpful discussions, and Yehuda Bergstein for insightful comments on the manuscript.

## Data and code availability

Behavioral data, processed eye-tracking data, and statistical analysis codes are shared on the OSF project page (https://osf.io/2v5xn/). Raw pre-processed data and pre-processing and features design codes are also available on the preregistered OSF project page.

## Declaration of Competing Interest

The authors filed a patent application US 17/954,409 “Automatic preference quantification of displayed objects based on eye tracker data” on 28 September 2022.

## Notes

### Competing Interest Statement

The authors have declared no competing interest.

