## Supplementary Material for "Passive identification of subjective preferences towards individual items using eye-tracking in a virtual reality environment"

**S1: Pilot study feature selection reassessment**

Feature selection reassessment on the pilot sample was done after the calculation correction, before the addition of Med_Scan_Speed features, to test whether the preregistered hypotheses still held. When the criteria described in the *feature selection* section were examined, we discovered that the Med_Dist feature was not one of the first features according to each of the criteria as was before the correction and therefore also preregistered (for example see features sorted in ascending order according to fixed effect p value in Table 1).

| | Feature | Fixed effect estimate | Fixed effect P value | Interaction estimate | Interaction P value |
|---|---|---|---|---|---|
| 1 | Max_Per_Dist | 1.50E-01 | 1.23E-03 | -1.51E-01 | 1.24E-02 |
| 2 | Med_diff_Dist | 1.45E-01 | 1.24E-03 | -2.06E-01 | 1.11E-04 |
| 3 | Min_Per_Diff_Dist | 1.57E-01 | 1.24E-03 | -2.04E-01 | 4.31E-04 |
| 4 | Max_Per_Diff_Obj_Dist | 7.37E-02 | 8.88E-03 | -1.12E-01 | 8.79E-02 |
| 5 | Max_Per_Diff_Dist | 6.74E-02 | 1.34E-02 | -1.58E-01 | 1.13E-02 |
| 6 | **Med_Dist** | 1.16E-01 | 1.34E-02 | -5.69E-02 | 3.50E-01 |
| 7 | Min_Per_Norm_Obj_Dist | -7.74E-02 | 5.74E-02 | 1.50E-01 | 6.84E-03 |
| 8 | Max_Per_Norm_Obj_Dist | -6.33E-02 | 8.83E-02 | 6.04E-02 | 2.55E-01 |
| 9 | Blinks_Sum_Constrained | 7.48E-02 | 9.01E-02 | -1.03E-01 | 5.69E-02 |
| 10 | Blinks_Num_Constrained | 6.31E-02 | 1.30E-01 | -8.15E-02 | 1.31E-01 |
| 11 | Blinks_Sum | 6.13E-02 | 1.73E-01 | -1.07E-01 | 5.18E-02 |
| 12 | Med_Norm_Obj_Dist | -4.62E-02 | 2.13E-01 | 1.06E-01 | 5.05E-02 |
| 13 | Blinks_Max_Constrained | 4.80E-02 | 2.79E-01 | -6.91E-02 | 1.97E-01 |
| 14 | Blinks_Num | 4.36E-02 | 2.87E-01 | -6.16E-02 | 2.53E-01 |
| 15 | Blinks_Mean_Constrained | 4.69E-02 | 2.92E-01 | -6.75E-02 | 2.04E-01 |
| 16 | Blinks_Mean | 4.89E-02 | 2.94E-01 | -8.40E-02 | 1.21E-01 |
| 17 | Blinks_Min_Constrained | 3.96E-02 | 3.43E-01 | -5.11E-02 | 3.27E-01 |
| 18 | Blinks_Min | 3.70E-02 | 3.85E-01 | -5.50E-02 | 2.97E-01 |
| 19 | GazedTime | -3.79E-02 | 3.86E-01 | 2.21E-02 | 6.92E-01 |
| 20 | Blinks_Max | 3.28E-02 | 4.78E-01 | -7.13E-02 | 1.89E-01 |
| 21 | Min_Per_Diff_Obj_Dist | 1.40E-02 | 7.13E-01 | 1.09E-02 | 8.48E-01 |
| 22 | Med_Diff_Obj_Dist | 8.21E-03 | 8.39E-01 | 5.37E-03 | 9.30E-01 |
| 23 | Min_Per_Dist | -9.33E-03 | 8.47E-01 | 7.23E-02 | 2.27E-01 |

**Table 1: Pilot sample features relation to preference ranking.** For each feature's mixed linear model: the fixed effect estimate, the fixed effect p value, the interaction estimate, and the interaction p value are shown. Features are sorted in ascending order according to the fixed effect p value, that was one of the selection criteria.

**S2: Features relation to ranking**

The relation of features to ranking was tested in mixed multilinear models adjusted for objects dimension. The results for valence and arousal ranking with features sorted in ascending order according to fixed effect p value are shown in Table 2 and 3 respectively.

| | Feature | Fixed effect estimate | Fixed effect P value | Interaction estimate | Interaction P value |
|---|---|---|---|---|---|
| 1 | **Med_Dist** | 2.94E-01 | 3.45E-21 | -2.44E-01 | 8.06E-12 |
| 2 | Max_Per_Dist | 2.54E-01 | 2.38E-20 | -2.20E-01 | 4.11E-10 |
| 3 | **Med_Scan_Speed** | 2.55E-01 | 1.31E-16 | -2.62E-01 | 1.30E-16 |
| 4 | Min_Per_Scan_Speed | 1.85E-01 | 4.12E-13 | -1.89E-01 | 4.08E-09 |
| 5 | Med_Diff_Obj_Dist | 1.65E-01 | 6.67E-13 | -1.68E-01 | 1.72E-07 |
| 6 | Med_diff_Dist | 1.53E-01 | 1.10E-09 | -1.81E-01 | 7.18E-09 |
| 7 | Max_Per_Norm_Obj_Dist | -1.12E-01 | 5.82E-08 | 1.29E-01 | 3.24E-05 |
| 8 | Max_Per_Diff_Dist | 1.18E-01 | 5.37E-07 | -1.46E-01 | 2.87E-06 |
| 9 | Max_Per_Scan_Speed | 1.03E-01 | 3.83E-06 | -9.93E-02 | 1.28E-03 |
| 10 | Min_Per_Dist | 1.31E-01 | 5.84E-06 | -8.19E-02 | 1.90E-02 |
| 11 | Min_Per_Diff_Dist | 1.10E-01 | 5.02E-05 | -1.31E-01 | 7.86E-05 |
| 12 | Min_Per_Norm_Obj_Dist | -8.47E-02 | 1.19E-04 | 1.29E-01 | 6.49E-05 |
| 13 | Blinks_Sum_Constrained | 9.73E-02 | 1.52E-04 | -1.28E-01 | 4.60E-05 |
| 14 | Max_Per_Diff_Obj_Dist | 7.40E-02 | 3.31E-04 | -1.22E-01 | 6.48E-05 |
| 15 | Blinks_Num | 8.26E-02 | 4.78E-04 | -1.02E-01 | 1.32E-03 |
| 16 | Blinks_Sum | 8.50E-02 | 6.99E-04 | -1.17E-01 | 2.40E-04 |
| 17 | Med_Norm_Obj_Dist | -6.74E-02 | 8.64E-04 | 1.06E-01 | 8.76E-04 |
| 18 | Blinks_Num_Constrained | 7.17E-02 | 3.45E-03 | -9.66E-02 | 2.40E-03 |
| 19 | Blinks_Max_Constrained | 7.26E-02 | 3.55E-03 | -1.00E-01 | 1.22E-03 |
| 20 | Blinks_Mean_Constrained | 6.70E-02 | 6.96E-03 | -9.81E-02 | 1.51E-03 |
| 21 | GazedTime | -7.06E-02 | 9.17E-03 | 7.59E-02 | 2.21E-02 |
| 22 | Blinks_Mean | 5.08E-02 | 3.89E-02 | -7.13E-02 | 2.18E-02 |
| 23 | Blinks_Max | 4.88E-02 | 4.45E-02 | -7.23E-02 | 2.09E-02 |
| 24 | Blinks_Min_Constrained | 3.28E-02 | 1.61E-01 | -5.91E-02 | 5.00E-02 |
| 25 | Min_Per_Diff_Obj_Dist | 3.04E-02 | 1.72E-01 | -3.55E-02 | 2.61E-01 |
| 26 | Blinks_Min | 2.05E-02 | 3.82E-01 | -4.20E-02 | 1.64E-01 |

**Table 2: Joined sample features relation to valence ranking.** For each feature's mixed linear model: the fixed effect estimate, the fixed effect p value, the interaction estimate, and the interaction p value are shown. Features are sorted in ascending order according to the fixed effect p value.

| | Feature | Fixed effect estimate | Fixed effect P value | Interaction estimate | Interaction P value |
|---|---|---|---|---|---|
| 1 | **Med_Dist** | -1.38E-01 | 4.93E-08 | 4.63E-02 | 1.35E-01 |
| 2 | Max_Per_Dist | -1.00E-01 | 2.07E-05 | -1.24E-02 | 6.85E-01 |
| 3 | **Med_Scan_Speed** | -8.91E-02 | 6.69E-04 | 5.05E-02 | 7.15E-02 |
| 4 | Min_Per_Scan_Speed | -6.37E-02 | 2.70E-03 | 4.63E-02 | 1.00E-01 |
| 5 | Min_Per_Dist | -7.44E-02 | 3.09E-03 | 4.38E-03 | 8.83E-01 |
| 6 | Med_Diff_Obj_Dist | -5.70E-02 | 3.83E-03 | 8.40E-02 | 2.54E-03 |
| 7 | GazedTime | 4.71E-02 | 5.00E-02 | 1.22E-02 | 6.71E-01 |
| 8 | Max_Per_Scan_Speed | -3.13E-02 | 1.03E-01 | -6.48E-02 | 1.59E-02 |
| 9 | Max_Per_Diff_Dist | -3.19E-02 | 1.07E-01 | -3.81E-02 | 1.64E-01 |
| 10 | Blinks_Num | -3.46E-02 | 1.16E-01 | 2.85E-03 | 9.17E-01 |
| 11 | Med_diff_Dist | -3.75E-02 | 1.24E-01 | -1.89E-03 | 9.45E-01 |
| 12 | Blinks_Sum | -3.13E-02 | 1.32E-01 | -2.55E-02 | 3.53E-01 |
| 13 | Blinks_Sum_Constrained | -2.97E-02 | 1.57E-01 | -1.99E-02 | 4.64E-01 |
| 14 | Blinks_Num_Constrained | -2.74E-02 | 1.77E-01 | -2.76E-02 | 3.13E-01 |
| 15 | Min_Per_Diff_Obj_Dist | -2.58E-02 | 1.98E-01 | 4.06E-02 | 1.39E-01 |
| 16 | Max_Per_Diff_Obj_Dist | 2.16E-02 | 2.15E-01 | 3.60E-02 | 1.66E-01 |
| 17 | Blinks_Max | -1.90E-02 | 3.63E-01 | -2.72E-02 | 3.15E-01 |
| 18 | Min_Per_Diff_Dist | -2.17E-02 | 3.70E-01 | -1.57E-02 | 5.84E-01 |
| 19 | Blinks_Max_Constrained | -1.85E-02 | 3.76E-01 | -2.11E-02 | 4.32E-01 |
| 20 | Min_Per_Norm_Obj_Dist | 1.75E-02 | 4.00E-01 | -4.14E-02 | 1.33E-01 |
| 21 | Blinks_Mean_Constrained | -1.49E-02 | 4.77E-01 | -1.49E-02 | 5.78E-01 |
| 22 | Max_Per_Norm_Obj_Dist | 1.35E-02 | 4.78E-01 | -2.56E-02 | 3.39E-01 |
| 23 | Blinks_Min_Constrained | -1.07E-02 | 5.95E-01 | -1.27E-02 | 6.26E-01 |
| 24 | Blinks_Min | 9.82E-03 | 6.24E-01 | -3.23E-02 | 2.15E-01 |
| 25 | Med_Norm_Obj_Dist | 5.19E-03 | 7.85E-01 | -2.96E-02 | 2.78E-01 |
| 26 | Blinks_Mean | -1.30E-03 | 9.50E-01 | -3.11E-02 | 2.46E-01 |

**Table 3: Joined sample features relation to arousal ranking.** For each feature's mixed linear model: the fixed effect estimate, the fixed effect p value, the interaction estimate, and the interaction p value are shown. Features are sorted in ascending order according to the fixed effect p value.

**S3: Pupil diameter features relation to ranking**

The relation of the pupil diameter features to ranking was also tested in mixed multilinear models adjusted for objects dimension. The results for preference, valence and arousal ranking with features sorted according to fixed effect p value are shown in Table 4-6 respectively.

| | Feature | Fixed effect estimate | Fixed effect P value | Interaction estimate | Interaction P value |
|---|---|---|---|---|---|
| 1 | Per50_PupilL | -3.13E-01 | 3.20E-13 | 2.90E-01 | 4.16E-11 |
| 2 | Per10_PupilL | -2.86E-01 | 4.28E-12 | 2.55E-01 | 5.47E-09 |
| 3 | Per50_PupilR | -2.65E-01 | 1.13E-10 | 2.42E-01 | 7.62E-09 |
| 4 | Per95_PupilL | -2.78E-01 | 1.80E-10 | 2.56E-01 | 1.53E-07 |
| 5 | Per10_PupilR | -2.51E-01 | 4.14E-10 | 2.23E-01 | 8.96E-08 |
| 6 | Per95_PupilR | -2.49E-01 | 2.18E-09 | 2.22E-01 | 2.95E-06 |

**Table 4: Joined sample pupil diameter features relation to preference ranking.** For each feature's mixed linear model: the fixed effect estimate, the fixed effect p value, the interaction estimate, and the interaction p value are shown. Features are sorted in ascending order according to the fixed effect p value.

| | Feature | Fixed effect estimate | Fixed effect P value | Interaction estimate | Interaction P value |
|---|---|---|---|---|---|
| 1 | Per50_PupilL | -4.96E-01 | 1.25E-20 | 3.53E-01 | 1.18E-13 |
| 2 | Per10_PupilL | -4.57E-01 | 1.20E-19 | 3.18E-01 | 1.68E-11 |
| 3 | Per50_PupilR | -4.38E-01 | 4.21E-18 | 3.05E-01 | 2.18E-11 |
| 4 | Per10_PupilR | -4.08E-01 | 7.65E-17 | 2.79E-01 | 6.03E-10 |
| 5 | Per95_PupilL | -4.14E-01 | 4.99E-16 | 2.86E-01 | 2.83E-08 |
| 6 | Per95_PupilR | -3.80E-01 | 2.13E-14 | 2.69E-01 | 6.10E-08 |

**Table 5: Joined sample pupil diameter features relation to valence ranking.** For each feature's mixed linear model: the fixed effect estimate, the fixed effect p value, the interaction estimate, and the interaction p value are shown. Features are sorted in ascending order according to the fixed effect p value.

| | Feature | Fixed effect estimate | Fixed effect P value | Interaction estimate | Interaction P value |
|---|---|---|---|---|---|
| 1 | Per50_PupilR | 1.11E-01 | 4.96E-03 | 7.15E-02 | 8.42E-02 |
| 2 | Per50_PupilL | 1.13E-01 | 5.60E-03 | 8.31E-02 | 5.41E-02 |
| 3 | Per10_PupilR | 1.09E-01 | 6.04E-03 | 6.13E-02 | 1.34E-01 |
| 4 | Per95_PupilL | 1.17E-01 | 9.21E-03 | 5.09E-02 | 2.89E-01 |
| 5 | Per10_PupilL | 1.02E-01 | 1.10E-02 | 8.26E-02 | 5.39E-02 |
| 6 | Per95_PupilR | 1.10E-01 | 1.49E-02 | 2.43E-02 | 5.95E-01 |

**Table 6: Joined sample pupil diameter features relation to arousal ranking.** For each feature's mixed linear model: the fixed effect estimate, the fixed effect p value, the interaction estimate, and the interaction p value are shown. Features are sorted in ascending order according to the fixed effect p value.

**S4: Med_Dist relation to ranking**

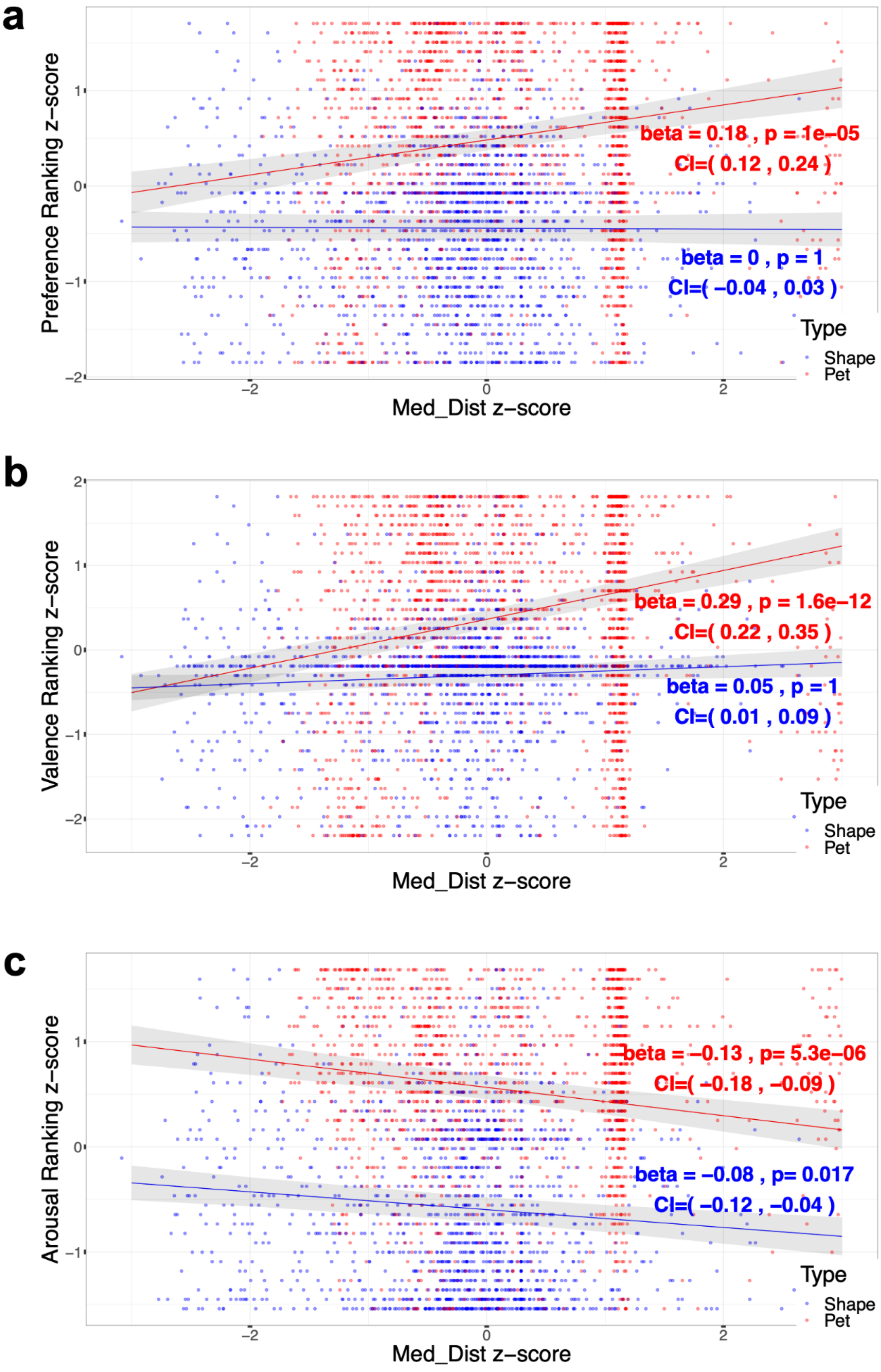
a
Preference Ranking z−score
beta = 0.18 , p = 1e−05
CI=( 0.12 , 0.24 )
beta = 0 , p = 1
CI=( −0.04 , 0.03 )
Type
Shape
Pet
Med_Dist z−score
b
Valence Ranking z−score
beta = 0.29 , p = 1.6e−12
CI=( 0.22 , 0.35 )
beta = 0.05 , p = 1
CI=( 0.01 , 0.09 )
Type
Shape
Pet
Med_Dist z−score
c
Arousal Ranking z−score
beta = −0.13 , p= 5.3e−06
CI=( −0.18 , −0.09 )
beta = −0.08 , p= 0.017
CI=( −0.12 , −0.04 )
Type
Shape
Pet
Med_Dist z−score

**Fig. 1**: **Med_Dist relation to Rankings.** Scatter plots of the subjective rankings as a function of Med_Dist per participant per object, and the linear models predicted values of rankings as a function of Med_Dist for each object type (pets in red and shapes in blue; fitted lines are surrounded by gray polygons of the 95% confidence interval). **a:** Preference ranking as a function of Med_Dist. A significant interaction of Type was obtained (beta = -0.18, $t[106.4] = -4.9$, $p = 2.6e-4$, CI = (-0.26,-0.11)) as described by a significant positive slope only for the pets in the plot. **b:** Valence ranking as a function of Med_ Dist. A significant interaction of Type was obtained (beta = -0.23, $t[222.2] = -6.2$, $p = 2e-7$, CI = (-0.31,-0.16)) as described by a significant positive slope only for the pets in the plot. **c:** Arousal ranking as a function of Med_ Dist. Type interaction was not found for arousal ranking (beta = 0.05, $t[606] = 1.6$, $p = 1$, CI = (-0.01,0.11)) as described by a significant negative slope of Med_Dist relation to arousal ranking for both types in the plot. (All variables are z-scored, and significance is noted for the Med_Dist estimate in a mixed multilinear model adjusted for objects dimension and percent time gaze for each type, where $p < 0.001$ '***', $p < 0.01$ '**', $p < 0.05$'*', $p < 0.1$ '.'; all tests are Bonferroni corrected for 3*32 ranking times the number of features = 96 tests).

**S5: Stimuli ranking density distributions**

The ranking density distribution across participants for the different ranking measures and object types are shown in the Fig. 2-7 that follows.

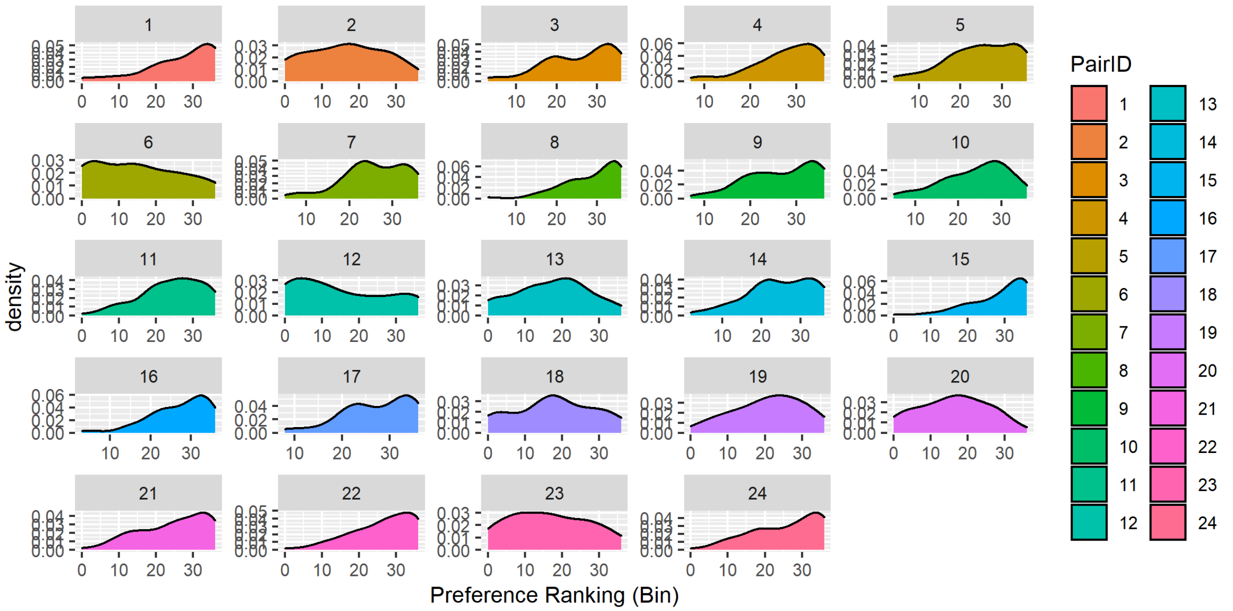


**Fig. 2: Pets preference density distribution.** Preference ranking density distribution across participants with valid preference ranking (N = 75). Each panel describes the ranking density distribution of a pet with a PairID and a corresponding color that matches to its control shape.

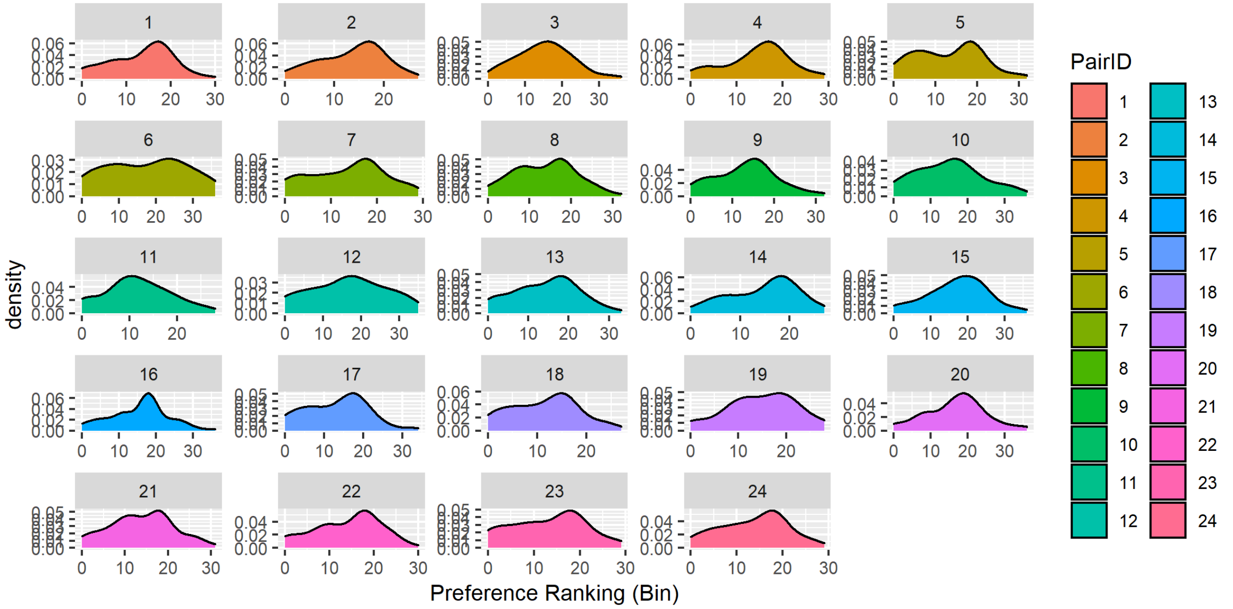


**Fig. 3: Shapes preference ranking density distribution.** Preference ranking density distribution across participants with valid preference ranking (N = 75). Each panel describes the ranking density distribution of a shape with a PairID and a corresponding color that matches to its control pet.

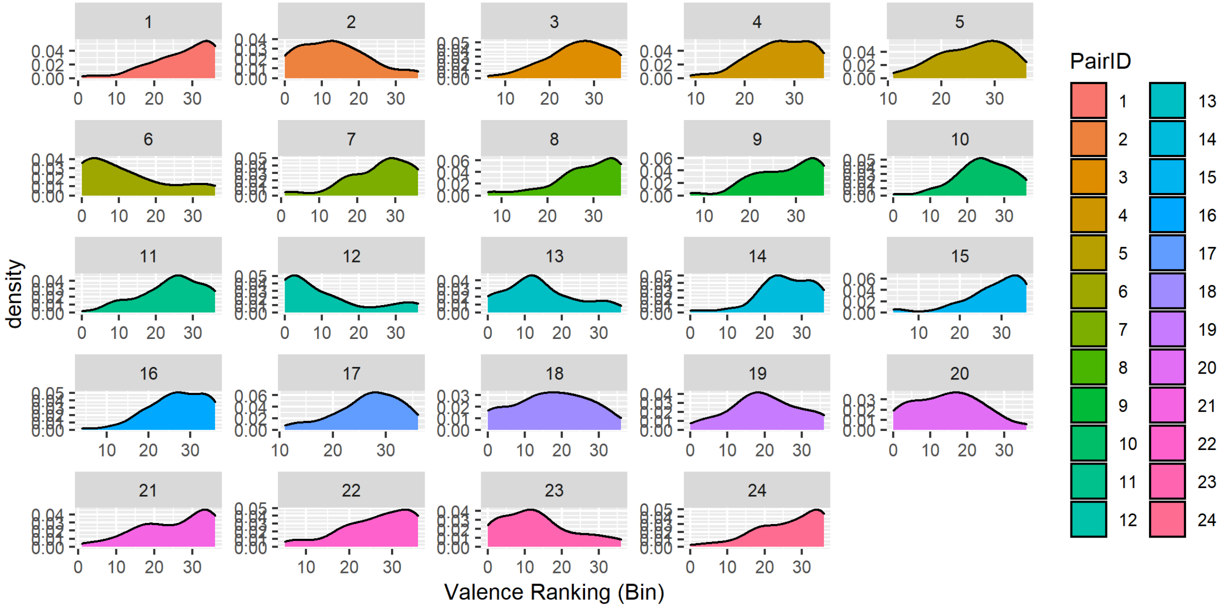


**Fig. 4: Pets valence ranking density distribution.** Valence ranking density distribution across participants with valid preference ranking (N = 77). Each panel describes the ranking density distribution of a pet with a PairID and a corresponding color that matches to its control shape.

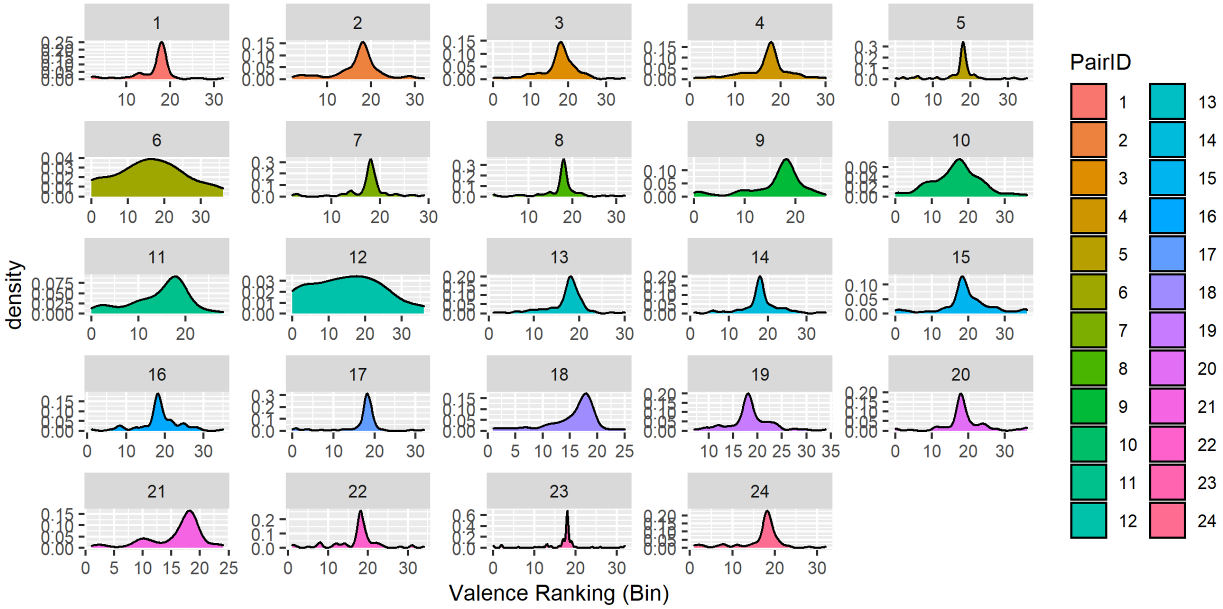


**Fig. 5: Shapes valence ranking density distribution.** Valence ranking density distribution across participants with valid preference ranking (N = 77). Each panel describes the ranking density distribution of a shape with a PairID and a corresponding color that matches to its control pet.

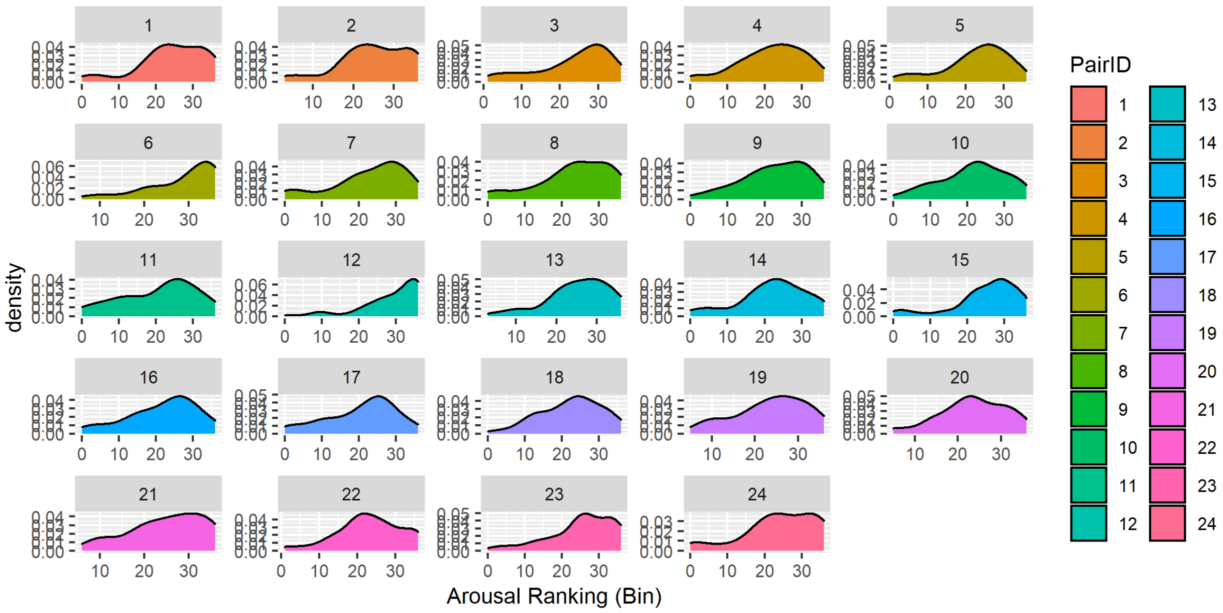


**Fig. 6: Pets arousal ranking density distribution.** Arousal ranking density distribution across participants with valid preference ranking (N = 66). Each panel describes the ranking density distribution of a pet with a PairID and a corresponding color that matches to its control shape.

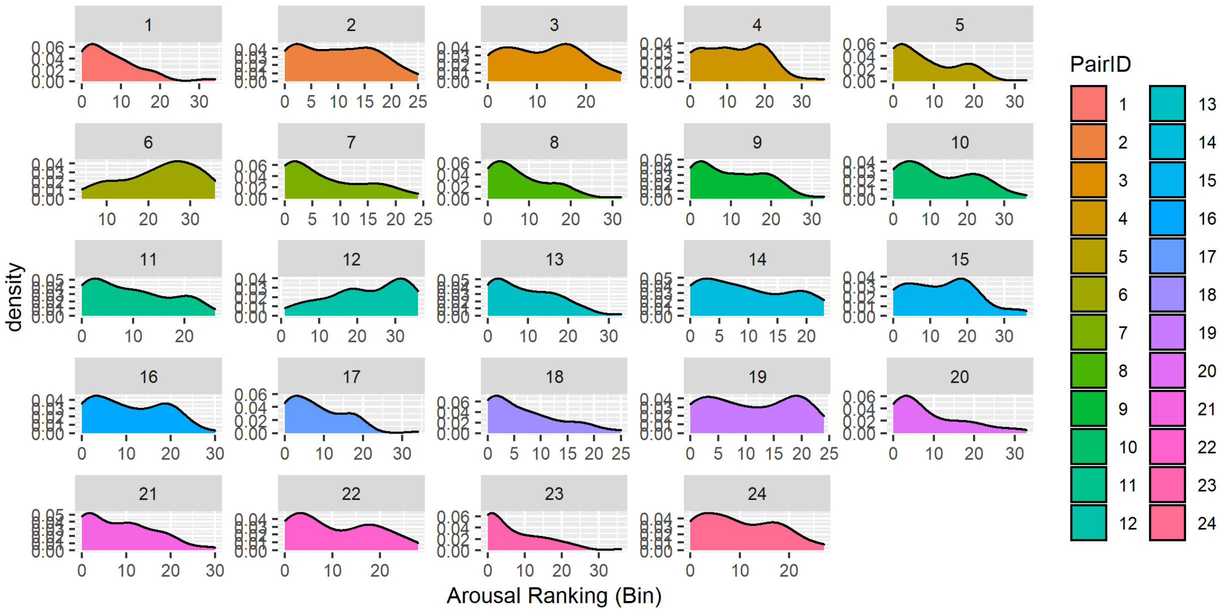


**Fig. 7: Shapes arousal ranking density distribution.** Arousal ranking density distribution across participants with valid preference ranking (N = 66). Each panel describes the ranking density distribution of a shape with a PairID and a corresponding color that matches to its control pet.
